# Elevated expression of complement C4 in the mouse prefrontal cortex causes schizophrenia-associated phenotypes

**DOI:** 10.1101/2020.11.27.400960

**Authors:** Mélanie Druart, Marika Nosten-Bertrand, Stefanie Poll, Sophie Crux, Felix Nebeling, Célia Delhaye, Yaëlle Dubois, Marion Leboyer, Ryad Tamouza, Martin Fuhrmann, Corentin Le Magueresse

**Affiliations:** INSERM UMR-S 1270, 75005 Paris, France; Sorbonne Université, 75005 Paris, France; Institut du Fer à Moulin, 17 rue du Fer à Moulin, 75005 Paris, France; Neuroimmunology and Imaging Group, German Center for Neurodegenerative Diseases, 53175 Bonn, Germany; INSERM U955, Neuro-Psychiatrie Translationnelle, Université Paris-Est, Créteil, France; AP-HP, DMU IMPACT, Département Médical Universitaire de Psychiatrie, Hôpitaux Universitaires Henri Mondor, Créteil, France

**Author notes:** Equal contribution (Marika Nosten-Bertrand, Stefanie Poll, Sophie Crux, Felix Nebeling).

## Abstract

Accumulating evidence supports immune involvement in the pathogenesis of schizophrenia, a severe psychiatric disorder. In particular, high expression variants of *C4*, a gene of the innate immune complement system, were shown to confer susceptibility to schizophrenia. However, how elevated C4 expression may impact brain circuits remains largely unknown. We used *in utero* electroporation to overexpress C4 in the mouse prefrontal cortex (PFC). We found reduced glutamatergic input to pyramidal cells of juvenile and adult, but not of newborn C4-overexpressing (C4-OE) mice, together with decreased spine density, which mirrors spine loss observed in the schizophrenic cortex. Using time-lapse two-photon imaging *in vivo*, we observed that these deficits were associated with decreased dendritic spine gain and elimination in juvenile C4-OE mice, which may reflect poor formation and/or stabilization of immature spines. In juvenile and adult C4-OE mice we found evidence for NMDA receptor hypofunction, another schizophrenia-associated phenotype, and synaptic accumulation of calcium-permeable AMPA receptors. Alterations in cortical GABAergic networks have been repeatedly associated with schizophrenia. We found that functional GABAergic transmission was reduced in C4-OE mice, in line with diminished GABA release probability from parvalbumin interneurons, lower GAD67 expression and decreased intrinsic excitability in parvalbumin interneurons. These cellular abnormalities were associated with working memory impairment. Our results substantiate the causal relationship between an immunogenetic risk factor and several distinct cortical endophenotypes of schizophrenia, and shed light on the underlying cellular mechanisms.

## Introduction

Genome-wide association studies (GWAS) identified genes involved in immune and in autoimmune processes that are associated with Schizophrenia (SZ)^1-4^. The major GWAS locus in SZ lies within the major histocompatibility complex (MHC) locus on chromosome 64. This includes the C4 gene which consists of C4A and C4B gene copy number variations and insertion/deletion of the human endogenous retrovirus K (HERVK). A detailed analysis of this locus showed a strong relationship between the genetic diversity of the C4 region and SZ^5^. In particular, the presence of several copies of C4A with HERVK insertions correlated with higher C4A expression and increased risk of SZ. The complement pathway is a central determinant of synaptic pruning, a developmental process in which the synaptic connections between neurons are refined in the brain from childhood until early adulthood^6^. Thus, the identification of C4A as a risk factor in SZ established a link between complement-mediated synaptic pruning and dendritic spine loss in the cortex of schizophrenic patients, a characteristic feature of the disease which had long been hypothesized to reflect excessive synapse refinement in adolescence^7, 8^. Moreover, C4A gene copy numbers within the human C4 locus are associated with increased neuronal complement deposition and microglial synapse uptake in co-cultures of human iPSC-derived neurons and microglial cells^9^.

In mice, the C4 gene presents homology with both the C4A and C4B genetic motifs of the human C4 gene locus. Connectivity between the retina and the visual thalamus is impaired in C4 knock-out mice^5^, indicating that the mouse C4 protein participates in synaptic pruning. It has been hypothesized that high C4 expression may result in increased synaptic pruning in upper layers of the neocortex^5^. Indeed C4A and C4B gene copy numbers positively correlated with neuropil contraction, which is often considered as a mesoscale correlate of developmental synapse elimination^10^. C4A gene copy numbers are also associated with increased neuronal complement deposition and microglial synapse uptake in co-cultures of human iPSC-derived neurons and microglial cells^9^. Furthermore, a recently published study showed that elevated C4 expression in the mouse prefrontal cortex causes dendritic spine loss, increased engulfment of synaptic material by microglial cells and impaired social behaviors^11^.

In addition to spine loss, several neural endophenotypes have been identified in the cortex of schizophrenic patients. Decreased NMDA receptor (NMDAR)-mediated neurotransmission in SZ has been inferred from at least four lines of evidence^12^: i) NMDAR antagonists induce schizophrenic-like behaviors; ii) the expression of NMDAR subunits is reduced in post-mortem brain tissue from schizophrenic patients iii) auto-antibodies against NMDA receptors cause clinical symptoms of SZ; iv) *GRIN2A*, an NMDAR subunit-encoding gene, has been associated with SZ in recent GWAS. Moreover, multiple deficits in GABAergic circuits have been identified in SZ. Abnormalities in GABAergic networks include reduced GABA concentration in the brain of schizophrenic patients, decreased expression of GAD67, an enzyme responsible for GABA synthesis, and abnormalities at parvalbumin-expressing (PV) neuron-pyramidal cell synapses^13^. What causes these distinct alterations, and how they relate to each other, is poorly understood. The relationship between C4 and SZ-associated cortical endophenotypes such as reduced NMDA receptor-mediated transmission, loss of GAD67 expression and altered GABAergic synapses remains unknown.

Here we examined the consequences of complement C4 overexpression in layer II/III pyramidal cells of the mouse prefrontal cortex on neuronal circuit development and function. C4 overexpression recapitulated several cortical phenotypes that have been associated SZ, including decreased spine number and reduced NMDA receptor function in pyramidal cells. Intriguingly, dendritic spine loss was accompanied by decreased spine turnover in juvenile mice. We also report several abnormalities in GABAergic circuits in C4-OE mice which are consistent with the alterations of GABAergic markers seen in the brain of SZ patients. Finally, our results reveal that the selective overexpression of C4 in PFC layer II/III is sufficient to impair working memory, which is reminiscent of SZ-associated cognitive deficits.

## Methods

A brief summary of experimental procedures is provided here with additional details available in the Supplemental Information accompanying this article. Additionally, Supplementary Tables S1-4 provide an overview of the number of recordings, animals and statistics in the various experiments.

### Animals

All animal experiments were performed according to guidelines of the European Community for the use of animals in research and were approved by the local ethical committees (C2EA-05 in France; LANUV NRW in Germany). All experiments were performed on mice that had been electroporated at embryonic stage E15 and were on a C57Bl/6J background.

### *In utero* electroporation

Pregnant mice at E15 were anesthetized by isoflurane 4% and were injected with Flunixin (0.05mg/kg body weight) before surgery. The surgery was performed under isoflurane 2% on a heated blanket and breathing was carefully monitored during all the procedure. Between 0.5 μl to 1μl of a solution containing pCAG-C4HA vector (0.8 μg/μl) together with pCAG-TdTomato vector (0.4 μg/μl) (molar ratio 4:1) in NaCl 0.9% were injected into the bilateral ventricles. In the control condition, a solution containing pCAG-TdTomato (0.8 μg/μl) alone was injected. Electrical pulses (50V; 50 ms) were applied five times at intervals of 950 ms with an electroporator (see Supplementary Information).

### *In vivo* two-photon imaging

Two-photon imaging was performed right after surgery at a Zeiss 7 multiphoton microscope with a Zeiss 20 x objective (NA = 1). TdTomato fluorescence was excited at 980 nm using an InSight X3 tunable laser (Spectra-Physics) and collected through a 617/73 band-pass filter on a highly sensitive non-descanned detector. After the first imaging session, the mice woke up and were placed in a cage with their littermates for 4-5 h (P15-18). For the second imaging session the mice were again anesthetized with isoflurane (initially 3%, then 1-1.5% for anesthesia maintenance) and imaged to retrieve the same dendritic structures recorded in the first imaging session (see Supplementary Information).

### Histology

Mice aged P30 were anesthetized with pentobarbital (150 mg/kg) and intracardially perfused with PBS followed by 4% buffered formaldehyde (Histofix, Roti). Free-floating sections were processed for immunohistochemistry as described in the Supplementary Information. Images were acquired using a confocal microscope (SP5, Leica) and analyzed according to procedures described in the Supplementary Information.

### Acute slice preparation, *ex vivo* electrophysiology and optogenetics

250 µm-thick slices were prepared from brains of control and C4-OE mice. For patch-clamp recordings, slices were transferred to the recording chamber where they were continuously superfused with ACSF (30-32°C) buffered by continuous bubbling with 95% O_2_-5% CO_2_. Whole-cell current and voltage-clamp recordings were performed in PFC layer II/III. Stimulus delivery and data acquisition were performed using Patchmaster software (Multichannel Systems). Signals were acquired with an EPC10-usb amplifier (Multichannel Systems), sampled at 20 kHz and filtered at 4 kHz. For optogenetic experiments, pulses of blue light (wavelength 470 nm) were emitted by a light emitting diode (CoolLED). Offline analysis was performed using Clampfit (Molecular Devices) and Igor Pro (Wavemetrics). For the study of miniature post synaptic currents, recordings were filtered offline at 2 kHz and analyzed using MiniAnalysis (Synaptosoft). See Supplementary Information.

### Behavior

Tests for locomotor activity (actimeter, Open Field), and working memory (learning of a Non-Matching-To-Sample task, Odor Span Test, Spatial Span Test) were performed and analyzed as described in the Supplementary Information.

### Statistical analysis

Data are presented as mean ± SEM. Statistical analyses were performed using Prism (Graphpad). The normality of data distribution was tested using Shapiro-Wilk’s test. Unpaired two-tailed T-tests (for normally distributed datasets) or Mann–Whitney tests (for non-normally distributed datasets) were used for comparisons between two groups. For multiple comparisons we used two-way ANOVA followed by Sidak’s test. Values of *P* < 0.05 were considered statistically significant. *P* values are reported as follows: * *P* < 0.05; ** *P* < 0.01; *** *P* < 0.001; **** *P* < 0.0001.

## Results

### Generation of C4-overexpressing (C4-OE) mice using *in utero* electroporation

Using in utero electroporation in C57Bl/6J mice we selectively overexpressed C4, the mouse homolog of the human *C4A* and *C4B* genes, in layer II/III of the prefrontal cortex (PFC) (Supplementary figure S1A). To identify C4-OE neurons using immunohistochemistry, an HA tag was added at the N-terminal end of the C4 construct. In addition, we co-electroporated the C4 construct with a tdTomato-expressing construct, in order to enable the identification of C4-expressing, tdTomato-positive (tdTom+) neurons in acute brain slices (Supplementary figure S1B-C). To ensure that most tdTom+ neurons indeed overexpress C4, we co-electroporated the C4 construct and the tdTomato construct with a 4:1 ratio. Immunohistochemistry for HA and TdTomato confirmed that 98.7 ± 1.8 % of tdTom+ neurons also express C4 (Supplementary figure S1D-E). Control mice were electroporated with the tdTomato construct only. We observed no neuronal mispositioning in C4-OE mice, suggesting the absence of major migration defect (Supplementary figure S2).

### Loss of excitatory synapses in C4-OE neurons from juvenile and adult, but not neonatal mice

We first evaluated passive and active cellular properties in tdTom+ layer II/III pyramidal cells using whole-cell patch-clamp *ex vivo*. The firing properties, membrane resistance, cell capacitance and resting membrane potential were similar in control and C4-OE neurons (Supplementary figure S3; Supplementary table S1). We next examined excitatory synaptic transmission at three developmental stages. The frequency of miniature excitatory postsynaptic currents (mEPSCs) in control tdTom+ cells markedly increased from early postnatal development (age P9-P10) to the juvenile stage (P25-P30), consistent with increased synapse generation, before decreasing from the juvenile stage to adulthood (P150-P220), in line with synapse elimination in the maturing cortex (Figure 1A-B; Supplementary table S1). The paired-pulse ratio of evoked EPSCs, a measure which is sensitive to alterations in presynaptic release probability, did not change significantly in control tdTom+ cells throughout development (Figure 1C-D; Supplementary table S1). Interestingly, mEPSC frequency was similar in control and C4-OE mice at P9-P10, but was significantly lower in C4-OE mice in juvenile and adult mice (Figure 1B; two-way ANOVA, *P* < 0.0001; juvenile: Sidak’s post-hoc test *P* < 0.0001; adulthood: Sidak’s post-hoc test, *P* < 0.05; Supplementary table S1). The paired-pulse ratio of evoked EPSCs did not differ between C4-OE and control mice in any age group (Figure 1D). We also observed a decrease in mEPSC amplitude from early development to the juvenile stage, but this was comparable in control and C4-OE mice (Figure 2B; Supplementary table S1).

**Figure 1.**
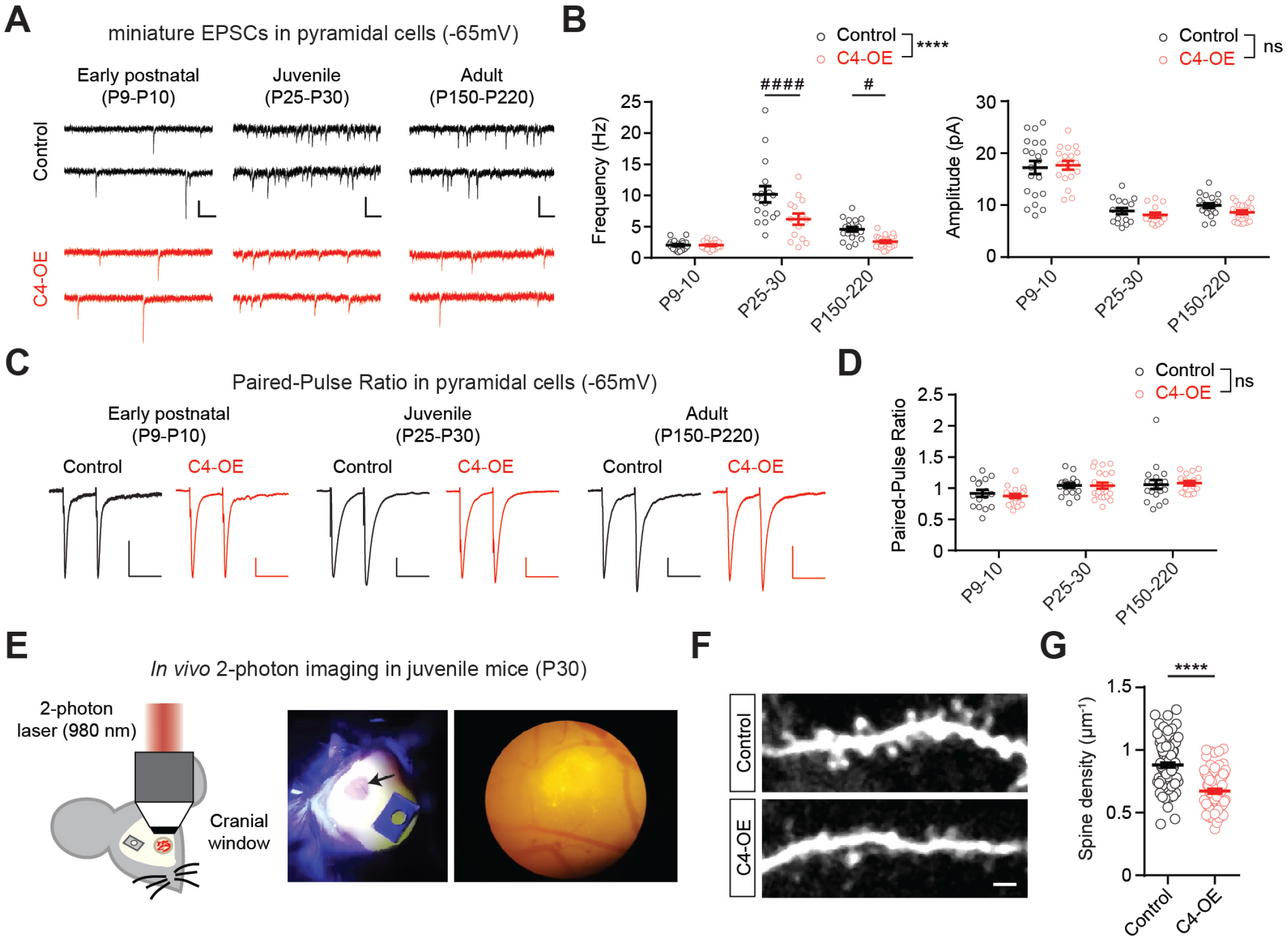
Reduced excitatory input onto C4-OE pyramidal cells. ***(A)*** Sample traces of mEPSCs in control and C4-OE pyramidal cells at three developmental stages; scale bars: 20pA/100ms. ***(B)*** mEPSC frequency was affected by C4-OE in juvenile and adult mice, but not in pups (two-way ANOVA, main effect, **** *P* < 0.0001; Sidak’s post-hoc test #### *P* < 0.0001, # *P* < 0.05, n.s. not significant). mEPSC amplitude was not significantly affected (control: early postnatal, n = 21 cells from 3 mice; juvenile, n = 16 cells from 5 mice; adult, n = 20 cells from 4 mice; C4-OE: early postnatal, n = 21 cells from 3 mice; juvenile, n = 16 cells from 5 mice; adult, n = 20 cells from 4 mice). ***(C)*** Average traces of eEPSCs evoked by paired-pulse stimulation with 50 ms interpulse interval, recorded from control and C4-OE pyramidal cells at three developmental stages; scale bars: 50 pA/50 ms. ***(D)*** Similar paired-pulse ratio in control and C4-OE pyramidal cells (n.s., not significant. control: early postnatal, n = 16 cells from 3 mice; juvenile, n = 17 cells from 3 mice; adult, n = 19 cells from 5 mice; C4-OE: early postnatal, n = 21 cells from 3 mice; juvenile, n = 21 cells from 3 mice; adult, n = 17 cells from 4 mice). All data are represented as mean ± SEM. Each open circle represents an individual cell. ***(E)*** Left, schematic drawing illustrating two-photon *in vivo* imaging in PFC at P30. Right, cranial window. ***(F)*** Examples of apical dendrites from electroporated pyramidal cells in control and C4-OE mice aged P30. ***(G)*** C4-OE decreased dendritic spine density in electroporated pyramidal cells from P30 mice; scale bar: 2 μm. (T-test, **** *P* < 0.0001; control: n = 99 dendrites from 10 mice; C4-OE: n = 76 dendrites from 7 mice). All data are represented as mean ± SEM. Each open circle represents an individual dendrite.

**Figure 2.**
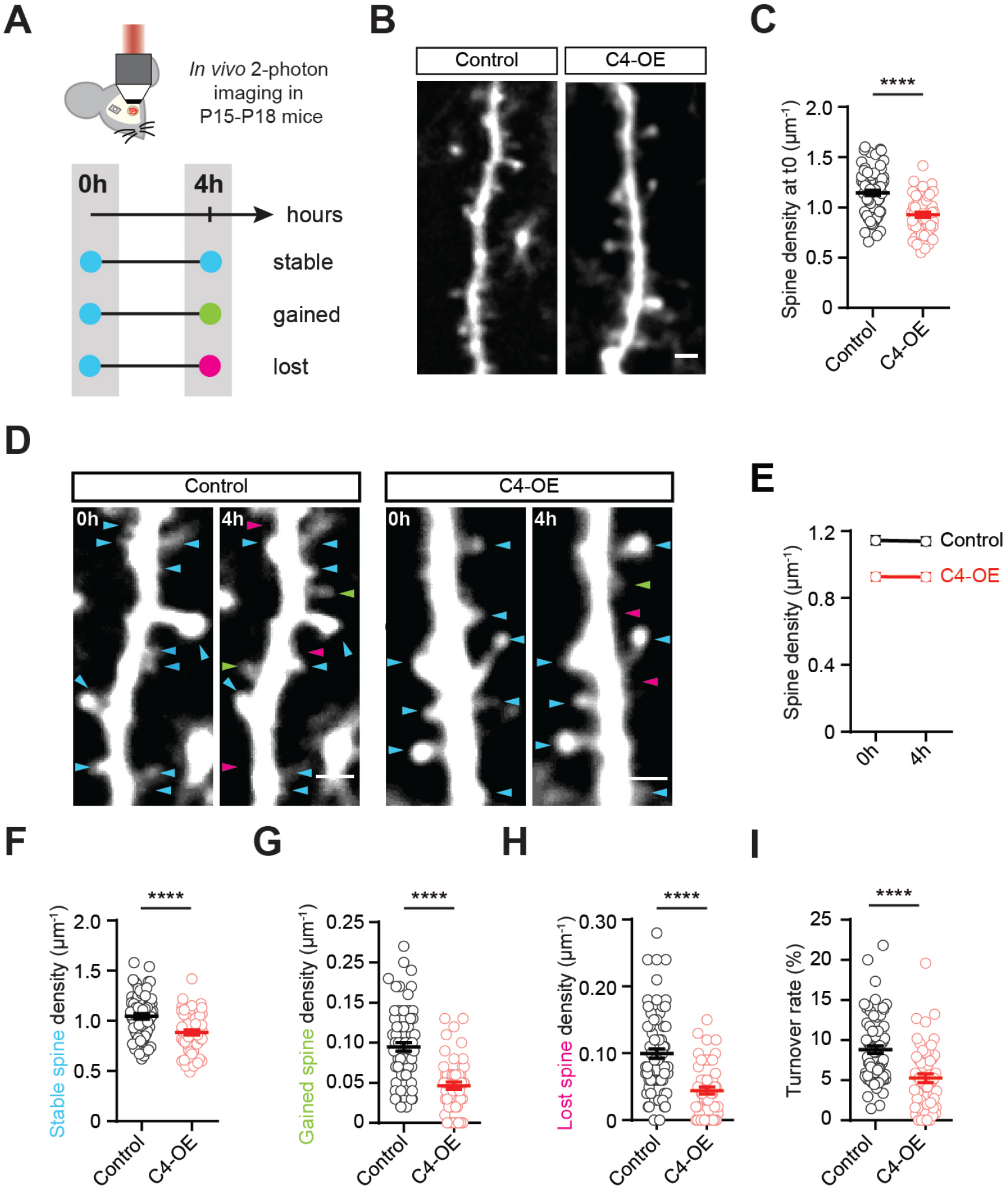
Decreased spine turnover in young (P15-P18) C4-OE mice. ***(A)*** Experimental timeline of *in vivo* two-photon imaging to investigate spine formation and elimination in electroporated pyramidal neurons. Spines already present at t_0_ are in blue; spines that were stable, gained or lost at t_0_ + 4h are in blue, green and magenta, respectively. ***(B)*** Examples of apical dendrites from control and C4-OE electroporated pyramidal neurons imaged at t_0_; scale bar: 2 μm. ***(C)*** Spine density at t_0_ in tdTom+ pyramidal neurons (t-test, **** *P* < 0.0001) ***(D)*** Example images showing spine turnover from t_0_ to t_0_ + 4h in control and C4-OE pyramidal cells; scale bar: 2 μm. ***(E)*** Evolution of the spine density between t_0_ and t_0_ + 4h in control and control and C4-OE pyramidal cells. ***(F-H)*** Stable, gained and lost spine density in control and C4-OE neurons. C4-OE caused a significant reduction in the number of both gained and lost spines (stable and gained spines: t-test, **** *P* < 0.0001; lost spines: Mann-Whitney test, **** *P* < 0.0001). ***(I)*** Spine turnover in control and C4-OE dendrites. The turnover rate is decreased in C4-OE dendrites (Mann-Whitney test, **** *P* < 0.0001). (Control: n = 73 dendrites from 5 mice; C4-OE: n = 55 dendrites from 7 mice). All data are represented as mean ± SEM. Each open circle represents an individual dendrite.

### C4 overexpression results in diminished formation and elimination of spines in juvenile PFC

To establish whether decreased functional excitatory input in C4-OE pyramidal cells correlates with reduced spine density, we imaged dendritic spines of tdTom+ pyramidal cells in layer I/II of the prefrontal cortex using *in vivo* 2-photon imaging in juvenile (P30) mice (Figure 1E). We found a substantial spine density decrease in C4-OE neurons (Figure 1F-G; T-test, *P* < 0.0001; Supplementary table S2). These results suggested that the reduction in spine density in C4-OE neurons is due to imbalanced spine dynamics (gain and loss) occurring prior to P30. Therefore, we performed dynamic imaging studies in mice aged P15-P18, which corresponds to a period of heightened structural synaptic plasticity^14-16^. Here, we imaged dendrites twice at a 4-hour interval (Figure 2A) to study fast spine kinetics at this stage of active synaptic reorganization. Similar to the P30 cohort, C4-OE neurons in P15-P18 mice displayed a robust decrease in spine density (Figure 2B-C; T-test, *P* < 0.0001; Supplementary table S2). While the overall spine density within the experimental groups did not change over the course of 4 h (Figure 2D-E; table S2), C4 overexpression had a strong effect on the structural plasticity of single spines; first, C4-OE neurons formed fewer stable spines (Figure 2F; T-test, *P* < 0.0001; Supplementary table S2). Second, C4-OE neurons showed decreased numbers of both gained and lost spines (Figure 2G-H; gained: T-test, *P* < 0.0001; lost: Mann-Whitney Test, *P* < 0.0001; also see Figure 2F; Supplementary table S2) resulting in an overall significantly reduced turnover of dendritic spines (Figure 2I; Mann-Whitney test, *P* < 0.0001; Supplementary table S2). Here we show that C4 overexpression results in diminished formation/stabilization and elimination of spines in juvenile PFC *in vivo*.

### NMDAR hypofunction in C4-OE pyramidal cells

To investigate NMDAR-mediated transmission in the cortex of C4-OE mice, we measured the AMPA/NMDA ratio of evoked EPSCs in tdTom+ cells. The AMPA/NMDA ratio was comparable in control and C4-OE neonates, but was significantly increased in juvenile and adult C4-OE mice (Figure 3A-B; Two-way ANOVA, *P* < 0.0001; juvenile and adult mice: Sidak’s test, *P* < 0.001; Supplementary table S1). The amplitude of mEPSCs, which is determined by the number of AMPAR per synapse, was not altered by C4 overexpression at any age examined (Figure 1B; Supplementary table S1). Furthermore, the decay time constant of NMDAR-mediated currents, which depends on the subunit composition of NMDA receptors, was similar in control and C4-OE mice of the same age group (Figure 3C; Supplementary table S1). Therefore the increased AMPA/NMDA ratio in C4-OE neurons results from reduced NMDAR-mediated currents. Increased AMPA/NMDA ratio has been repeatedly associated with altered synaptic expression of the AMPAR GluA2 subunit^17, 18^ that controls the calcium permeability of AMPARs^19, 20^. Therefore we assessed the rectification of synaptic AMPAR-mediated currents which is sensitive to the presence of the GluA2 subunit. We found that the rectification index of evoked AMPAR currents was significantly decreased in juvenile and adult C4-OE mice, but not in pups (Figure 3D-E; two-way ANOVA, *P* < 0.0001; juvenile: Sidak’s test, *P* < 0.001; adulthood: Sidak’s test, *P* < 0.05; Supplementary table S1). This suggests that synaptic AMPARs in C4-OE neurons show reduced GluA2 subunit content, thus becoming more calcium-permeable, concomitantly with NMDAR hypofunction.

**Figure 3.**
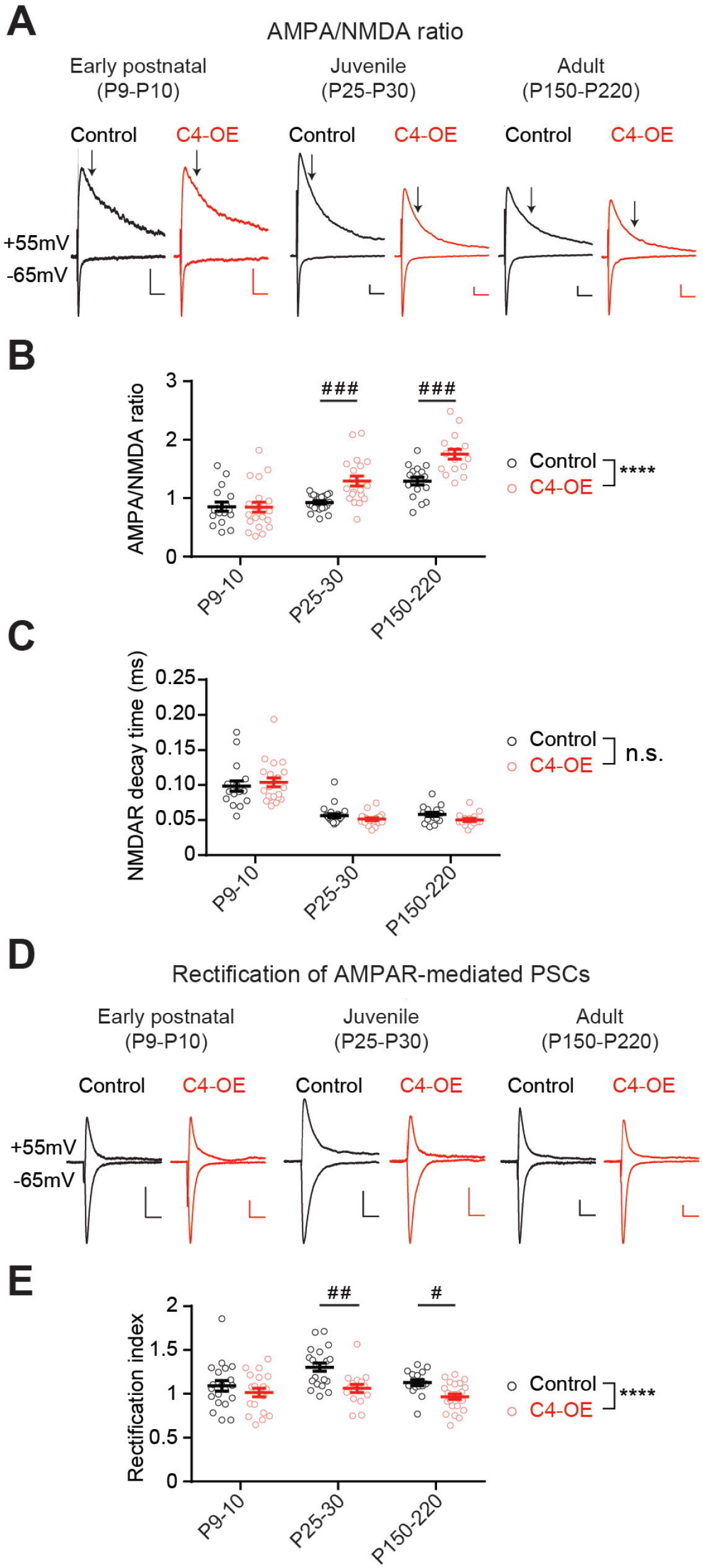
Altered AMPA/NMDA ratio and AMPAR rectification index in C4-OE mice. ***(A)*** Sample traces of evoked synaptic AMPA and NMDA currents recorded at −65 mV and +55 mV respectively in the presence of GABAzine at 3 developmental stages; The NMDAR component was measured 50 ms after stimulation (arrows). The AMPAR component was measured as the peak of the evoked EPSC at −65mV. Scale bars: 50pA/50ms. ***(B-C)*** AMPA/NMDA ratio and NMDA decay time throughout development in control and C4-OE pyramidal cells. The AMPA/NMDA ratio was increased in C4-OE neurons from juvenile and adult mice, but not from neonates. The decay time of NMDARs was not affected by C4-OE at any stage of development, indicating that C4-OE does not alter NMDAR subunit composition. This together with unchanged mEPSC amplitude in C4-OE mice (Fig.2B) indicates that the increased AMPA/NMDA ratio reflects reduced NMDAR-mediated excitatory transmission in C4-OE pyramidal neurons (two-way ANOVA, main effect, **** *P* < 0.0001, Sidak’s post-hoc test ### *P* < 0.001; n.s., not significant. Control: early postnatal, n = 17 cells from 3 mice; juvenile, n = 23 cells from 4 mice; adult, n = 17 cells from 4 mice; C4-OE: early postnatal, n = 21 cells from 3 mice; juvenile, n = 20 cells from 5 mice; adult, n = 16 cells from 4 mice). ***(D)*** Sample traces of evoked AMPA-mediated currents recorded at −65mV and +55mV at 3 developmental stages; scale bars: 50 pA/50 ms. ***(E)*** The rectification index of AMPR-mediated currents was decreased in pyramidal cells from juvenile and adult C4-OE mice, suggesting the insertion of GluA2 subunit-lacking AMPAR at excitatory synapses (two-way ANOVA, **** *P* < 0.0001, Sidak’s post-hoc test ## *P* < 0.01, # *P* < 0.05. Control: early postnatal, n = 20 cells from 3 mice; juvenile, n = 21 cells from 3 mice; adult, n = 17 cells from 4 mice; C4-OE: early postnatal, n = 20 cells from 3 mice; juvenile, n = 16 cells from 3 mice; adult, n = 24 cells from 3 mice). All data are represented as mean ± SEM. Open circles represent single cells.

### C4 overexpression results in reduced GABAergic transmission

Abnormalities in GABAergic networks constitute another hallmark of the schizophrenic cortex. They include synaptic abnormalities as well as decreased expression of GAD67 in parvalbumin-expressing (PV) interneurons. We examined spontaneous inhibitory synaptic transmission in tdTom+ pyramidal cells from juvenile mice using a high chloride intracellular solution to maximize the signal/noise ratio. Both the frequency and amplitude of miniature inhibitory postsynaptic currents (mIPSCs) were significantly decreased in C4-OE neurons (Figure 4A-B; frequency: Mann-Whitney test, *P* = 0.0004; amplitude: T-test, *P* = 0.0003; Supplementary table S1). Reduced mIPSC frequency may result from decreased release probability from GABAergic terminals in the maternal immune activation (MIA) model of SZ^21^ which has been associated with increased expression of complement C1q and C4^22, 23^. We investigated GABA release probability using a similar protocol^21^. IPSCs were evoked by four consecutive electrical stimuli delivered at 20 Hz while holding voltage-clamped tdTom+ pyramidal cells at a membrane potential of +10 mV (Figure 4C). Short-term synaptic depression, which is sensitive to changes in release probability, showed a non-significant trend towards reduction in C4-OE neurons (Figure 4D; RM two-way ANOVA, *P* = 0.089; Supplementary table S1). In the MIA model, altered release probability from PV neurons, but not from other types of interneurons, has been inferred from optogenetic studies of short-term plasticity^21^. We reasoned that selectively activating the PV neuron-pyramidal cells synapse in C4-OE mice might allow to uncover significant differences in short-term depression. To test this hypothesis, we expressed Channelrhodopsin2 (ChR2) fused to YFP in PV-expressing interneurons using timed matings between lox-STOP-lox ChR2 mice and PV-Cre mice (Supplementary figure S4A). Pregnant females were electroporated at embryonic stage E15 either with a tdTomato-encoding plasmid (control group) or with tdTomato + C4 (C4-OE condition). To verify the specificity of ChR2 expression, we patched-clamped ChR2-YFP+ cells in the control group and recorded their firing pattern in response to a 500 ms pulse of blue light (470 nm). All tested ChR2-YFP cells exhibited the characteristic fast-spiking firing pattern of PV interneurons (n = 3; Supplementary figure S4B). Light-evoked PSCs in pyramidal cells were blocked by GABAzine, confirming their GABAergic nature (Supplementary figure S4C-D). Next we assessed short-term depression of PV neuron IPSCs in pyramidal cells using four consecutive blue light pulses (0.5 ms) delivered at 20 Hz (Figure 4E). We found that PV neuron-pyramidal cell synapses in C4-OE mice displayed significantly reduced short-term depression, consistent with decreased GABA release probability (Figure 4F; two-way ANOVA repeated measures, main effect, *P* = 0.014; Supplementary table S1). Thus, similar to the MIA model, decreased release probability from PV neurons may contribute to diminish GABAergic transmission onto pyramidal cells in C4-OE mice.

**Figure 4.**
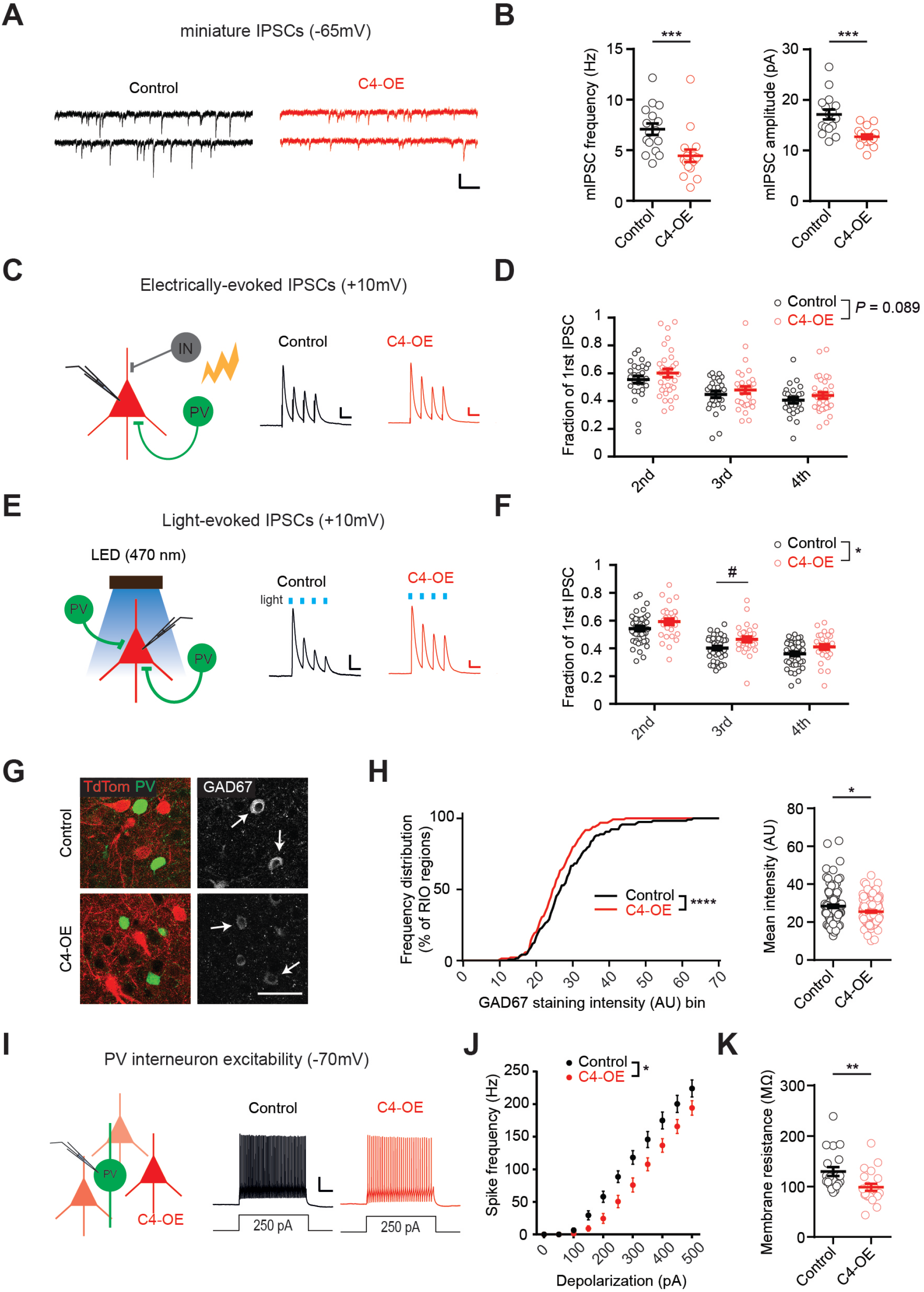
Alterations of GABAergic networks in C4-OE mice. ***(A)*** Sample traces of mIPSCs recordings at a holding potential of −65 mV in control and C4-OE pyramidal cells from juvenile mice (P25-P30); scale bars: 20pA/100ms. ***(B)*** Both mIPSC frequency and amplitude were decreased in C4-OE pyramidal cells (frequency: Mann-Whitney test *** *P* = 0.0004; amplitude: t-Test *** *P* = 0.0003. Control: n = 16 cells from 4 mice; C4-OE: n = 16 cells from 4 mice). ***(C)*** Schematic drawing illustrating the electrical stimulation of GABAergic release onto electroporated pyramidal cells (left) and example traces of 4 consecutive eIPSCs evoked with 50ms interpulse intervals, in control and C4-OE pyramidal cells (right); scale bars: 50 pA/50 ms. ***(D)*** Ratios of the second, third and fourth eIPSC peak amplitude to the first eIPSC peak amplitude. Paired-pulse depression exhibited a trend towards being decreased in C4-OE neurons (two-way RM ANOVA, main effect, *P* = 0.089. Control: n = 29 cells from 3 mice; C4-OE: n = 37 cells from 3 mice). ***(E)*** Schematic drawing illustrating the optogenetic activation of PV neuron and resulting IPSC recorded in electroporated pyramidal cells (left); and example traces of 4 consecutive light-evoked IPSCs with 50 ms interpulse intervals in control and C4-OE pyramidal cells (right); scale bars: 50 pA/50 ms. ***(F)*** Ratios of the second, third and fourth light-evoked IPSC peak amplitude to the first IPSC peak amplitude. The increased ratio in C4-OE pyramidal cells indicates reduced presynaptic GABA release at PV neuron-pyramidal cell synapses (two-way RM ANOVA, main effect, * *P* = 0.014, Sidak’s post-hoc test # *P* < 0.05. Control: n = 43 cells from 4 mice; C4-OE: n = 30 cells from 3 mice). All data are represented as mean ± SEM. Open circles represent single cells. ***(G)*** Example of immunostainings for tdTomato/PV and GAD67 on PFC slices from control and C4-OE mice. Arrows indicate GAD67-expressing PV cells; scale bar: 20µm. ***(H)*** Frequency distribution and mean ± SEM of GAD67 expression in PV+ cells. GAD67 expression in PV cells was decreased in C4-OE mice. Data points are averages from transfected region ROIs. (Frequency distribution: Kolmogorov-Smirnov test, **** *P* < 0.0001; Average GAD67 expression: Mann-Whitney test, * *P* = 0.011. Control: n = 117 ROI from 6 mice; C4-OE: 134 ROI from 6 mice). ROI, region of interest. ***(I)*** Schematic drawing depicting the patch-clamp recording of a PV interneuron located in the vicinity of tdTomato pyramidal cells (left) and sample spike trains in PV interneurons evoked by a 250 pA somatic current injection in control and C4-OE mice in the presence of synaptic blockers (right); scale bars: 20 mV/100 ms. ***(J)*** PV interneurons exhibited reduced intrinsic excitability in C4-OE mice (two-way RM ANOVA, main effect, * *P* = 0.013. Control: n = 20 cells from 3 mice; C4-OE: n = 21 cells from 4 mice). ***(K)*** The input resistance of PV neurons was significantly decreased in C4-OE mice. (Mann-Whitney test, ** *P* = 0.0024. Control: n = 20 cells from 3 mice; C4-OE: n = 21 cells from 4 mice).

### C4 overexpression in pyramidal cells causes reduced input resistance, decreased excitability, and lower GAD67 expression in PV neurons

The availability of presynaptic GABA is one of the factors determining release probability. GAD67 is an enzyme responsible for basal cortical GABA synthesis^24^, and reduced expression of GAD67, especially in PV cells, has been found in post-mortem cortical tissue from schizophrenic patients in multiple studies^13, 25^. In mouse models, reducing GAD67 expression in PV neurons decreases both the frequency of spontaneous inhibitory currents^26^ and the amplitude of evoked GABAergic currents in pyramidal cells^27^. Therefore we measured GAD67 expression in PV interneurons located in the vicinity of control or C4-OE tdTom+ pyramidal cells. As GAD67 is expressed in both presynaptic terminals and in the cell soma, we quantified GAD67 expression in the clearly delineated cell soma (Figure 4G)^21^. GAD67 expression in PV cells was significantly lower in C4-OE mice than in control mice (Figure 4H; frequency distribution, Kolmogorov-Smirnov test, *P* < 0.0001; Mean intensity, Mann-Whitney test, *P* = 0.011; Supplementary table S2). These results suggest that reduced GABA synthesis may contribute to the deficit in GABAergic transmission at PV neuron – pyramidal cell synapses in C4-OE mice. It has been proposed that reduced expression of activity-dependent biomarkers, including GAD67, in PV neurons, may result from decreased PV neuron activity in the schizophrenic cortex^13^. We investigated the number and distribution of PV neurons in the cortex of C4-OE juvenile mice and found that they were not altered (Supplementary figure S5; Supplementary table S2). We then electroporated C4 + tdTomato or tdTomato alone in PV-Cre::RCE mice which express EGFP selectively in PV neurons. We patch-clamped EGFP+ PV neurons located in the vicinity of tdTom+ pyramidal cells, in control and C4-OE mice aged P25-P30 (Figure 4I). The intrinsic excitability of PV neurons, i.e. their propensity to fire action potentials when subjected to an input current in the presence of drugs blocking synaptic transmission, was significantly diminished in the C4-OE condition (Figure 4J; RM two-way ANOVA, main effect *P* = 0.013; Supplementary table S3). This was associated with lower input resistance of PV cells (Figure 4K; Mann-Whitney test, *P* = 0.0024; Supplementary table S3), in the absence of significant changes in cell capacitance or resting membrane potential (Figure S6A-B; Supplementary table S3).

### C4 overexpression results in increased excitatory input on PV neurons

Since synaptic inputs also contribute to neuronal excitability, we recorded miniature excitatory and inhibitory inputs (mEPSCs and mIPSCs) in EGFP+ PV neurons. We observed a significant increase in mEPSC frequency, and a trend towards increased mEPSC amplitude, in the C4-OE condition (Supplementary figure S6C-D; frequency: Mann-Whitney test, *P* = 0.018; amplitude: Mann-Whitney test, *P* = 0.069; Supplementary table S3). The paired-pulse ratio of evoked EPSCs in PV neurons remained unchanged. (Figure S6E-F; table S3). Neither the frequency nor the amplitude of mIPSCs was changed following C4 overexpression in pyramidal cells (Supplementary figure S6G-H; Supplementary table S3). Thus, in contrast to C4-OE pyramidal cells, PV neurons exhibit increased excitatory input in C4-OE mice.

### Working memory deficits in C4-OE mice

Finally we investigated whether the changes that we observed at the cellular level may result in long-lasting behavioral alterations in C4-OE mice. In SZ, deficits in working memory are central to cognitive impairments and appear to result, at least in part, from abnormalities in PFC function^28, 29^. Therefore we asked whether increased C4 expression in the PFC alters working memory. Since classical working memory tests rely on locomotion, we first verified that control and C4-OE mice showed comparable spontaneous locomotor activity in an actimeter (Figure 5A; Supplementary table S4). Spontaneous activity of C4-OE mice also did not differ from control mice in the open field assay (Figure 5B; Supplementary table S4). We then used two different tests of working memory, namely the odor span test and the spatial span test. C4-OE mice performed equally well as control mice during the acquisition of the non-matching to sample task, in which mice learned to associate the presence of a new odor and food reward (Figure 5C-E; Supplementary table S4). In the subsequent span tests, however, when mice had to remember an increasing number of odors or positions to obtain the reward, C4-OE mice exhibited reduced working memory capacity compared to control mice (Figure 5F-I; odor span: Mann-Whitney test, *P* = 0.0022; spatial span: Mann-Whitney test, *P* = 0.045; Supplementary table S4). Taken together, these results indicate that working memory is impaired in C4-OE mice, in line with neuronal abnormalities in PFC networks.

**Figure 5.**
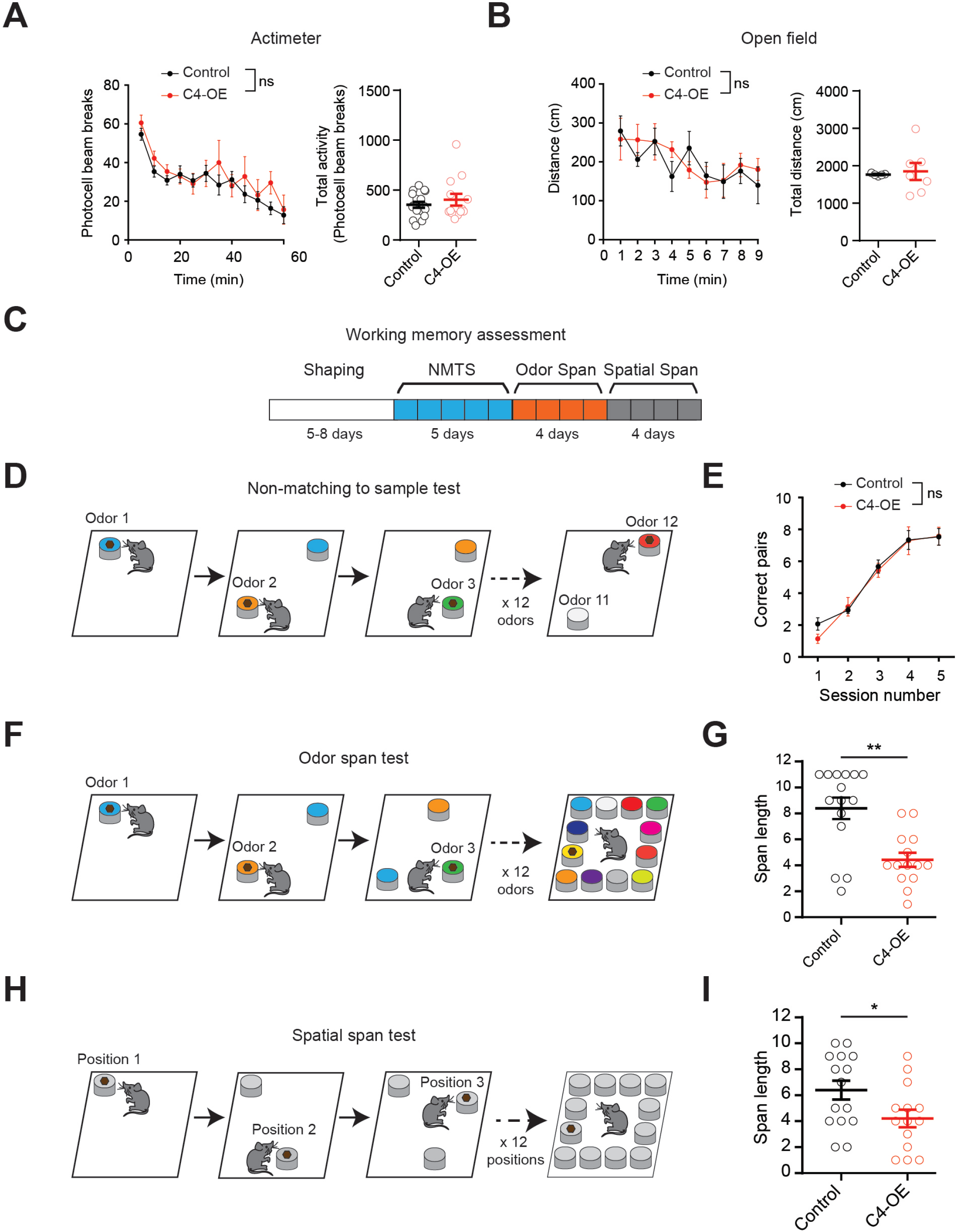
Impaired working memory in C4-OE mice. ***(A)*** Locomotor activity measured at weaning in an actimeter, in control (n = 17 mice) and C4-OE male mice (n = 13 mice). Each dot represents spontaneous activity averaged over a period of 5 minutes. ***(B)*** Locomotor activity measured one week after weaning in the open field, in control (n = 5 mice) and C4-OE mice (n = 7 mice). ***(C)*** Time line of working memory tests. Animals were first trained to dig into scented sawdust to find a reward pellet (shaping) before learning a non-matching to sample (NMTS) task, followed by 4 days of odor span test, and 4 days of spatial span test. ***(D)*** NMTS test procedure. Mice were exposed to pairs of odors and learned to associate a new odor with a reward. ***(E)*** Performance in the NMTS odor test was similar in control and C4-OE mice (Control: n = 15 mice; C4-OE: 14 mice). ***(F)*** Mice were exposed to an increasing number of odors in the odor span test. ***(G)*** Working memory capacity in the odor span test is impaired in C4-OE mice (Control: n = 15 mice; C4-OE: 14 mice; Mann-Whitney test, ** *P* = 0.0022). ***(H)*** Mice were exposed to an increasing number of positions in the spatial span test. ***(I)*** Working memory capacity in the spatial span test is also affected in C4-OE mice (Control: n = 15 mice; C4-OE: 14 mice; Mann-Whitney test, * *P* = 0.0448). All data are represented as mean ± SEM. Each open circle represents an individual mouse.

## Discussion

GWAS have provided unprecedented identification of predisposing genetic risk factors in SZ. High expression variants of the C4 gene contribute to the strong association between SZ and a genomic region within the MHC locus on Chromosome 6 in GWAS^4, 5^. Here we show that overexpressing C4 in the mouse PFC recapitulates several cortical phenotypes associated with SZ, including synapse loss, decreased NMDAR-mediated neurotransmission, impaired function of GABAergic circuits, and deficits in working memory.

### Decreased spine turnover in C4-OE mice

Elevated expression of C4 in neurons enhances microglia-mediated synaptic engulfment, and results in the selective elimination of small or medium-sized spines^9, 11^. Our *in vivo* imaging results clearly show an overall decrease in gained and lost dendritic spines in juvenile mice as a consequence of C4 overexpression. This apparent discrepancy may reflect an underappreciated role of complement in regulating spine formation or immature spine stabilization. Only the smallest spines are lost in cortical layer III in subjects with schizophrenia while larger spines are retained, which has led to the hypothesis that the formation or stabilization of immature synapses is affected, rather than the removal of established synapses^30^. Similarly, the decrease in spine turnover observed in our model could be due to excessive microglial phagocytosis of the synapses in formation or just formed, preventing their stabilization. The disappearance of these spines in less than 4 hours, the time window used for our time-lapse 2-photon microscopy acquisitions, would thus result in decreased spine gain. In this scenario, the more mature spines, which are the least likely to be eliminated, would constitute a higher proportion of the remaining spines in C4-OE neurons, resulting in reduced spine turnover.

It is also noteworthy that the decrease in excitatory input to C4-OE neurons is not present at the early stages of postnatal development, but only appears between P10 and P25, a period of strong synaptic remodeling in the cortex. This may be relevant for the pathophysiology of SZ, a disorder in which even the earliest prodromal symptoms occur at a relatively late stage of postnatal development.

### GABAergic network abnormalities

We show that C4-OE in pyramidal cells decreases GABA release from PV neurons onto pyramidal cells, consistent with decreased GAD67 expression in PV neurons and reduced PV neuron excitability. The cause of the well-substantiated decrease in GAD67 cortical expression in SZ is unclear. It has been hypothesized that lower activity of layer II/III pyramidal cells, possibly due to their reduced number of dendritic spines, could result in fewer excitatory inputs onto PV cells (for instance through lower neuronal activity-regulated pentraxin secretion from pyramidal cells), leading to reduced PV neuron excitation and hence decreased activity-dependent GAD67 expression^13, 31^. In C4-OE mice, reduced GAD67 expression in PV neurons may result from altered intrinsic properties of PV neurons, rather than reduced synaptic excitatory input. Indeed we show that PV neuron excitability is impaired in C4-OE mice, while increased mEPSC frequency in PV neurons indicates that PV neurons are spared from excitatory synapse loss associated with C4 overexpression in pyramidal cells.

### Linking neural dysfunctions and cognitive impairment in C4-OE mice

Cognitive dysfunction is a central feature of SZ that is not treated by current pharmacotherapy, and determines the functional outcome of the illness^32^. Impaired working memory has been proposed to be central in the cognitive impairments associated with SZ^28, 29^. Spine loss, reduced NMDAR transmission and PV neuron dysfunction in the PFC may all contribute to impair working memory. For example, decreased cortical connectivity in SZ has repeatedly been associated with working memory deficits^33^. Decreased synapse density in layer II/III may particularly affect connectivity, since layer II/III pyramidal cells make major contributions to cortico-cortical connections, including callosal connections, and therefore integrate information across cortical areas and hemispheres. Furthermore, blocking NMDA, but not AMPA, receptors during a working memory task abolishes persistent activity in prefrontal neurons^34^. Finally, PV neurons contribute to the mechanisms underlying working memory^35^, in line with the well-established role of PV neurons in the generation of fast, gamma frequency oscillations in neocortex^36, 37^. Our results suggest that these distinct alterations occur together during development as a consequence of elevated C4 expression, making it difficult to attribute cognitive dysfunction to a single one of them. The combination of reduced connectivity, NMDAR hypofunction and altered PV neuron function might collectively modify the dynamics of specific attractors, which are thought to be the substrate for persistent neural activity underlying short-term memory^38, 39^.

## Conclusion

Decreased spine density, reduced NMDAR function, and alterations in GABAergic networks have been associated with SZ. We show that elevated C4 expression in PFC pyramidal cells results in functional alterations that are consistent with these SZ-associated phenotypes, and causes working memory impairment, a characteristic feature of cognitive deficits in SZ. Future studies using transgenic mice with altered C4 expression in other brain regions or neuronal types may allow to further investigate the relationship between C4 and distinct phenotypes associated with SZ. In particular, we propose that the persistent decrease in NMDAR-mediated neurotransmission in C4-OE mice might be the consequence of impaired stabilization of immature synapses. This could provide a conceptual framework to relate the hypotheses of excessive synaptic pruning and NMDAR hypofunction in SZ.

## Acknowledgements

MD was the recipient of fellowships from Sorbonne University and from FRM/Venite Cantemus (FDT20190400802). The project was supported by funds from the Investissements d’Avenir program under reference ANR-11-IDEX-0004-02 to CLM and RT, the Emergence program of Sorbonne University to CLM, and from an ERANET Neuron Grant to RT (ANR-18-0008-01), MF (BMBF 01EW1905) and CLM (ANR-18-0008-02). We thank Drs L Maroteaux, P Gaspar and J-C Poncer for critically reading the manuscript.

