## Supplementary information for "Elevated expression of complement C4 in the mouse prefrontal cortex causes schizophrenia-associated phenotypes"

#### Materials and methods

##### Animals

All animal experiments were performed according to guidelines of the European Community for the use of animals in research and were approved by the local ethical committees (C2EA-05 in France; LANUV NRW in Germany). All experiments were performed on mice that had been electroporated at embryonic stage E15. Mice were housed in their home cages and kept under a 12-h light/dark cycle at 22 ± 2°C. Food and water were available ad libitum. For optogenetics experiments, we used electroporated transgenic Pvalb$^{tm1(cre)Arbr}$ $^{(+/-)}$::ChR2(H134R)-YFP$^{(lox/-)}$ mice obtained after crossing homozygous Pvalb$^{tm1(cre)Arbr}$ (JAX n°017320) females expressing Cre recombinase under the control of the Pvalb promoter (kindly provided by Dr JC Poncer, Institut du Fer à Moulin, Paris) and homozygous lox-STOP-lox ChR2-YFP males (JAX n°012569) (kindly provided by Dr MC Angulo, Institute of Psychiatry and Neuroscience, Paris). To identify and patch-clamp PV interneurons ex vivo, we used PV-Cre mice crossed with RCE:LoxP reporter mice (JAX n°010701) expressing enhanced green fluorescent protein (EGFP) under the control of the endogenous Gt(ROSA)26Sor promoter/enhancer regions and the CAG promoter. All mice were on a C57Bl/6J background. Females and males were used indifferently for electrophysiology, immunohistochemistry and imaging experiments. Only males were used for behavioral experiments.

##### DNA plasmid constructs

pCAG-HAC4 expression vector design was based on pCAG-HT2B-HA. The pCAG-HT2B-HA was a gift from I Moutkine (Fer à Moulin Institute, Paris). The HT2B gene was removed and replaced by the full coding sequence for murine C4b (NM_009780.2). The cDNA of C4b was obtained by reverse transcription PCR of total RNA from whole brain of adult 129S1/SvImJ mice. An HA tag was introduced at the N-terminus of the C4b sequence to allow the identification of the overexpressed C4 peptide. pCAG-tdTomato expression vector design was based on pCAG-mGFP which was a gift from Dr. C. Cepko (Matsuda and Cepko,

2004) (Addgene plasmid # 14757). mGFP was replaced by tdTomato in the pCAG backbone. Subcloning of C4b and tdTomato sequences were done using InFusi (Clonetech) and DNA were purified with Nucleospin kit (Macherey-Nagel). All constructs were verified by sequencing the entire coding region of the inserts.

.

#### *In utero* electroporation

Pregnant mice at E15 were anesthetized by isoflurane 4% and were injected with Flunixin (0.05mg/kg body weight) before surgery. The surgery was performed under isoflurane 2% on a heated blanket and breathing was carefully monitored during all the procedure.

A midline laparotomy was performed and the uterine horns were exposed and moistened with sterile NaCl 0.9%. Between 0.5µl to 1µl of a solution containing pCAG-C4HA vector (0.8 µg/µl) together with pCAG-TdTomato vector (0.4 µg/µl) (molar ratio 4:1) in NaCl 0.9% were injected into the bilateral ventricles with a glass micropipette made from a microcapillary tube (#5-000-1001-X10, Drummond Scientific). In the control condition, a solution containing pCAG-TdTomato (0.8 µg/µl) alone was injected. The injected plasmid solution contained Fast Green (0.1%) to monitor the injection. The embryo's head in the uterus was held between the tweezers-type electrode consisting of two disc electrodes of 5 mm diameter (CUY650P5, Nepagene). The cathode was placed just above the olfactory bulbs and the anode was placed at the back of the head below the cerebellum. Electrical pulses (50V; 50 ms) were applied five times at intervals of 950 ms with an electroporator (CUY21 Edit, Nepagene). The uterine horn was placed back into the abdominal cavity. The abdominal cavity was filled up with sterile NaCl 0.9% and the abdominal muscle and skin were sutured separately. After recovery, pregnant mice were returned to their home cage and sutures were exanimated every day until birth. Correct tdTomato expression was first checked through the intact skin shortly after birth (age P1-P2) and confirmed *post mortem* on brain slices from each experimental animal. Mice without tdTomato expression in the prefrontal cortex (PFC) were excluded from the analysis.

#### Cranial window surgery

At postnatal day 15 to 18 (P15-18) (and P30 respectively) a cranial window surgery was performed as previously described (Bittner et al., 2010; Fuhrmann et al., 2010; Fuhrmann et al., 2007). Mice were anesthetized with an intraperitoneal injection of ketamin/xylazin (0.13/0.01 mg/g body weight). For analgesia an additional subcutaneous injection of buprenorphin (0.05 mg/kg) was given. The anesthetic profile was deepened and maintained by inhalation of isoflurane (0.2-0.4% in 0.2 l/min oxygen). After the withdrawl reflex of the hind limb had subsided, the surgery begun. The eyes were covered with eye ointment (Bepanthen) to prevent drying and the mouse was placed on a heating pad to maintain body temperature. Subsequently, mice were fixed to a stereotactic frame and the skin of the head was disinfected with 70% ethanol. The skin above the skull was removed using surgical scissors. To assure correct placement of the cranial window, the expression of the fluorophor was located using a UV light source. Next, the skull bone was dried and a 4-mm-diameter circular piece of bone was removed over the prefrontal cortex using a dental drill. The dura mater was carefully removed and a circular coverslip (Ø 4 mm) was inserted into the craniotomy and glued to the bone. To allow stable and repetitive fixation of the mouse under the microscope, a small metal bar was attached next to the cortical window using UV-curable dental adhesive.

**Two-photon *in vivo* imaging and analysis**

Two-photon imaging was performed right after surgery at a Zeiss 7 multiphoton microscope with a Zeiss 20 x objective (NA = 1). The anesthesia was maintained with 1-1.5% inhalation of isoflurane while the mouse was head fixed to a custom-made imaging frame. TdTomato fluorescence was excited at 980 nm using an InSight X3 tunable laser (Spectra-Physics) and collected through a 617/73 band-pass filter on a highly sensitive non-descanned detector. First, a 600 µm (x) x 600 µm (y) x 400 µm (z) overview z-stack (4 µm z-spacing) was acquired to retrieve the region of interest over multiple sessions. Then z-stacks of single dendrites (1 µm z-spacing) with a pixel size of 0.089 µm were acquired. After the first imaging session, the mice woke up and were placed in a cage with their littermates for 4-5 h (P15-18). For the second imaging session the mice were again anesthetized with isoflurane (initially 3%, then 1-1.5% for anesthesia maintenance) and imaged to retrieve the same dendritic structures recorded in the first imaging session.

Datasets were blinded prior to analysis by an experimenter not involved in the image analysis. In each animal 4-19 dendrites of 30-120 µm length were analyzed. Spines that extended out laterally from the dendritic shaft were counted manually by scrolling through the z-stack, as previously described (Gu et al., 2014). Spine density was calculated by the number of spines divided by the length of the respective dendrite. Turnover of dendritic spines was calculated as follows: turnover [%] = 100*((#gained + #lost)/(#T1+#T2)). Here, #gained refers to the number of gained and #lost to the number of lost spines, respectively and #T1 and #T2 to the number of all spines at time-points T1 and T2, respectively. T1 and T2 refer to 0h and 4h.

**Histology**

*Immunohistochemistry*

Mice aged P30 were anesthetized with pentobarbital (150 mg/kg) and intracardially perfused with PBS followed by 4% buffered formaldehyde (Histofix, Roti). The brains were removed and fixed overnight in 4% Histofix. 40 µm sagittal sections were cut using a vibratome (VT1000S, Leica). Free-floating sections were permeabilized and blocked in 0.2% Triton-PBS, 3% BSA-PBS for 60 min before incubation with the first antibody in 0.2% Triton, 3% BSA-PBS overnight at 4°C. After three washes in PBS, sections were incubated with the secondary antibody in PBS for 2h, washed again in PBS, and mounted using Fluoromount G (Invitrogen). Primary antibodies were: rat anti-HA (1:300, Sigma), rabbit anti-Parvalbumin (PV27, 1:1000, Swant), mouse anti-Glutamate Decarboxylase 67 (GAD67, clone 1G10.2, 1:500, Millipore-Merk). Secondary antibodies were: goat anti-rat Alexa Fluor 488 (1:400, Invitrogen), goat anti-rabbit Cy5 (1:1000, Jackson Immuno Research), donkey anti-rabbit Cy3 (1:1000, Jackson Immuno Research), goat anti-mouse Alexa Fluor 647 (1:1000, Jackson Immuno Research). Hoechst 33342 was used for nuclear staining. Images were acquired using a confocal microscope (SP5, Leica).

*PV cell number and distribution*

The distribution of PV neurons in different PFC layers was quantified using maximal intensity projections of 3D confocal stacks. For each mouse, 4 PFC sagittal sections were imaged using a confocal microscope (Leica SP5, 40x objective). Layers were identified based on Hoechst nuclear staining and delineated using ImageJ's ROI manager software. The distribution of PV neurons in different PFC layers was quantified using maximal intensity projections of 3D confocal stacks. The density of PV cells (number of PV cells per surface unit) in each layer was obtained by averaging the density across the 4 slices. During image acquisitions and quantifications, the investigator was blind to the genotype.

*GAD67 expression*

GAD67 expression levels in PV cells were measured by analyzing confocal images (Leica SP5, 40x objective) using ImageJ. For each animal, 3 to 4 PFC sagittal slices were acquired. At least 80 PV neurons per mice were analyzed. A single confocal section located 10 µm below the surface of the slice was acquired in each slice. The same acquisition parameters were used for each slice and were set such that the brightness of the GAD67 immunostaining was below saturation values. The brightness of the PV immunostaining was set above saturation level, allowing to use PV as a marker to delineate the shape of PV neurons. The density of GAD67 immunostaining within PV cells was then automatically calculated using Image J.

During image acquisitions and quantifications, the investigator was blind to the condition (control or C4-OE).

**Acute slice preparation**

250 µm-thick coronal slices were prepared from brains of control and C4-OE mice. Mice were deeply anesthetized with sodium pentobarbital (150 mg/kg) and perfused transcardially with ice-cold (0–4°C) oxygenated (95% $O_2$-5% $CO_2$) solution containing (in mM): 110 Choline chloride, 2.5 KCl, 25 glucose, 25 NaHCO3, 1.25 NaH2PO4, 0.5 CaCl2, 7 MgCl2, 11.6 L-ascorbic acid and 3.1 sodium pyruvate. The brain was extracted and acute sagittal slices were cut in the same ice-cold solution using a vibroslicer (HM 650 V, Microm), then stored in artificial cerebrospinal fluid (ACSF) containing the following (in mM): 125 NaCl, 2.5 KCl, 25 glucose, 25 NaHCO3, 1.25 NaH2PO4, 2 CaCl2, and 1 MgCl2, continuously bubbled with 95% $O_2$-5% $CO_2$. Slices were incubated in ACSF at 32°C for 20 minutes and then at room temperature (20-25°C). For patch-clamp recordings, slices were transferred to the recording chamber where they were continuously superfused with ACSF (30-32°C) buffered by continuous bubbling with 95% $O_2$-5% $CO_2$.

**Electrophysiology and optogenetics**

Whole-cell current and voltage-clamp recordings were performed in PFC layer II/III. Recorded pyramidal cells all expressed tdTomato. Recorded PV interneurons were in the vicinity of electroporated tdTomato+ pyramidal cells and were visually identified based on the expression of EGFP (in PV-Cre::RCE mice) or YFP (in PV-Cre::lox-STOP-lox ChR2-YFP mice). Stimulus delivery and data acquisition were performed using Patchmaster software (Multichannel Systems). Signals were acquired with an EPC10-usb amplifier (Multichannel Systems), sampled at 20 kHz and filtered at 4 kHz. Offline analysis was performed using Clampfit (Molecular Devices) and Igor Pro (Wavemetrics). For the study of miniature post synaptic currents, recordings were filtered offline at 2 kHz and analyzed using MiniAnalysis (Synaptosoft).

Patch-clamp pipettes (3–6 Mohm resistance) were prepared from borosilicate glass (BF150-86-10; Harvard Apparatus) using a DMZ pipette puller (Zeitz). Current-clamp experiments were performed using the following intracellular solution (in mM): 105 K-gluconate, 10 HEPES, 10 phosphocreatine-Na, 0.3 Na3GTP, 4 MgATP, 30 KCl (pH 7.25, adjusted with KOH). For voltage-clamp experiments the following intracellular solution was used (in mM): 120 Cs-methane sulfonate, 10 CsCl, 10 Hepes, 10 Phosphocreatine, 0.2 EGTA, 8 NaCl, 2 ATP-Mg, 3 QX 314 (pH 7.25, adjusted with CsOH). To record miniature and evoked inhibitory postsynaptic currents (mIPSCs ans eIPSCs), we used an intracellular solution containing (in mM): 60 Cs methane sulfonate, 70 CsCl, 10 Hepes, 10 Phosphocreatine, 0.2 EGTA, 8 NaCl, 2 ATP-Mg (pH 7.25, adjusted with CsOH). Liquid junction potentials (-14.9 mV for Cs-based intracellular solutions and -5mV for the K+-based intracellular solution) were left uncorrected.

*Excitability*

Pyramidal cell and PV interneuron intrinsic properties and excitability were recorded in current-clamp in presence of the following synaptic blockers: SR95531 hydrobromide (Gabazine, 10 μM, Hello Bio), 6-cyano-7-nitroquinoxaline-2,3-dione (CNQX, 10 μM, Hello Bio) and D-2-amino-5-phosphonopentanoic acid (D-APV, 50 μM, Hello Bio). Spike frequency-depolarization curves were generated by injecting series of 500 ms depolarizing current steps increasing by 50 pA. The firing pattern of patched-clamped YFP+ cells in the cortex of PV-Cre::lox-STOP-lox ChR2-YFP mice was monitored using 500 ms pulse of blue light emitted by a light emitting diode (CoolLED).

*Miniature EPSCs and evoked EPSCs*

Miniature excitatory post synaptic currents (mEPSCs) were recorded at a holding potential of -65 mV in the presence of tetrodotoxin (TTX, 0.5 μM, Hello Bio) and Gabazine (10 μM). At least 100s were analysed for each recording. Evoked excitatory post synaptic currents (eEPSCs) were recorded at a holding potential of -65 mV and triggered at a frequency of 0.1 Hz in the presence of GABAzine (10 μM) with a monopolar electrode connected to a constant current stimulator (Iso-Flex, A.M.P.I.), positioned in the vicinity of the recorded neuron. At least 12 consecutive sweeps were acquired for each cell. The paired-pulse ratio (PPR) was calculated as the ratio of the peak amplitude of the averaged current response evoked by the second stimulation to the peak amplitude of the averaged current response evoked by the first stimulation.

*Miniature IPSCs and evoked IPSCs*

mIPSCs were recorded at a holding potential of -65 mV in the presence of TTX (0.5 μM, Hello Bio), CNQX, (10 μM) and D-APV (50 μM). At least 100 s were analysed for each recording. Evoked IPSCs were recorded at a holding potential of +10mV to avoid unclamped action potentials commonly associated with evoked IPSCs measured at hyperpolarized potentials in our high-chloride recording conditions, in the presence of CNQX (10 μM) and APV (50 μM). At least 25 consecutive sweeps were acquired and averaged for each cell. For electrically evoked IPSC (eIPSCs), four stimulations at 20 Hz were applied with a monopolar electrode connected to a constant current stimulator positioned in layer 3, 100-150 μm away from of the recorded neuron. For light-evoked IPSCs, the recorded pyramidal cell was placed in the center of the field of view. The response to four 0.5-ms pulses of blue light (470nm) at 20 Hz emitted by a light emitting diode (CoolLED) was recorded. The paired-pulse ratio of eIPSCs was calculated as the ratio of the peak amplitude of the averaged current response evoked by the second, third and fourth stimulations to the peak amplitude of the averaged current response evoked by the first stimulation.

*AMPA/NMDA ratio and rectification index of AMPAR-mediated currents*

To measure the AMPA/NMDA ratio and the rectification index, EPSC were evoked with a monopolar electrode positioned in layer 3, 100-150 μm away from the recorded neuron. At least 12 consecutive sweeps were acquired and averaged at each holding potential. The AMPA/NMDA ratio was recorded in presence of GABAzine (10 μM). The AMPA current ($I_{AMPA}$) was measured as the peak amplitude of the current response at a holding potential of -65mV. The NMDA current ($I_{NMDA}$) was measured at a holding potential of +55mV, 50ms after stimulation, when AMPAR-mediated current have receded due to the fast deactivation kinetics of AMPARs. The ratio is calculated as $I_{AMPA}/I_{NMDA}$.

To study the rectification AMPA-mediated currents, pyramidal cells were recorded in the presence of GABAzine (10 μM) and D-APV (50 μM). Spermine (10 μM) was added to the intracellular solution. The rectification index was defined as the peak amplitude of synaptic currents measured at a holding potential of +55 mV (≈ +40 mV after correcting for junction potential) multiplied by 2 and divided by the peak amplitude of synaptic currents measured at a holding potential of -65mV (≈ -80 mV after correcting for junction potential).

#### Behavior

*Actimeter*

We used actimeter racks of 8 circular corridors (Imetronic, http://www.imetronic.com/devices/circular-corridor/) equipped with infrared sensors to detect and automatically record locomotion. Animals were introduced individually in corridors and their spontaneous, novelty-driven activity was recorded for a total of 60 minutes. Activity, measured as photocell beam breaks, was averaged over periods of 5 minutes. Control and C4-OE male mice were tested at weaning (age P30).

*Open Field*

The test took place in a white open-field apparatus (ViewPoint, Lyon, France) under high illumination (~100 lux). Animal were individually placed in the corner of the open-field and were allowed to explore for 9 min. A video tracking system, which included a computer-linked overhead camera, was used to monitor general lon comotor activity (https://viewpoint.fr/en/p/software/videotrack, ViewPoint, Lyon, France). Control and C4-OE male mice were tested a week after weaning (age P35 – P39).

*Odor Span and Spatial Span Test*

Working memory capacity was tested using a protocol adapted from a previous study (Young et al., 2007). All scented mixtures were prepared with 100 g of sawdust, 18 ground powdered pellets (Chocapic, Nestle) to mask the scent of the reward pellet and 3 g of a ground powder odor (Savory, Cinnamon, Cumin, Nutmeg, Mint, Fennel, Four spices, Herbs of Provence, Coriander, Basil, Coffee, Ginger). During shaping and testing, adult (4- to 6-month-old) male mice were maintained at 80-85% of their body weight. During the shaping phase, mice were trained to dig into a bowl of unscented sawdust to retrieve the hidden reward pellet. Once all animals had learned the procedure, the non-matching to sample (NMTS) task started for 5 consecutive days. Each day, the mouse was placed on a square platform and required to dig into the first bowl (odor 1) to retrieve the reward and to remember the scent of that bowl. Upon consumption of the reward pellet, a second baited bowl (odor 2) was added. The first bowl (no longer baited) and the new bowl were randomly distributed on the platform. Following retrieval of the pellet, the first bowl was removed and a novel baited bowl (odor 3) randomly placed. This process was repeated until each individual mouse had been exposed to 11 pairs of scented bowls or stopped after a maximum of 12 minutes when the mouse did not perform the task, and the number of correct pairs (direct digging into the baited bowl) was recorded.

Following the NMTS, animals were then tested 4 consecutive days for the odor span task, using the same procedure except that, following reward retrieval, the no longer baited bowls were not removed so that mice had to remember an increasing numbers of odors (span). The span length was calculated as the number of correct responses with more than one bowl on the platform prior to the first mistake. Thus, using 12 odors, the maximum span length is 11. The span score was given by the first mistake, but each mouse was allowed to continue the task until the 12 odors had been sampled or for a maximum of 12 minutes. The span length was the best score obtained by each tested mouse over a period of 4 consecutive days.

The same mice were subsequently tested for 4 consecutive days in the spatial span task. They had to remember an increasing numbers of locations, following a procedure similar to the odor span test, except that bowls of unscented sawdust were identified according to their spatial location on the platform. The span length was the best score obtained by each tested mouse over a period of 4 consecutive days.

The investigator was blind to the condition (control or C4-OE) during behavioral tests and analysis.

**Statistical analysis**

Data are presented as mean ± SEM. Statistical analyses were performed using Prism (Graphpad). The normality of data distribution was tested using Shapiro-Wilk's test. Unpaired two-tailed T-tests (for normally distributed datasets) or Mann–Whitney tests (for non-normally distributed datasets) were used for comparisons between two groups. For multiple comparisons we used two-way ANOVA followed by Sidak's test. Values of $P < 0.05$ were considered statistically significant. $P$ values are reported as follows: * $P < 0.05$; ** $P < 0.01$; *** $P < 0.001$; **** $P < 0.0001$.

**Resource table**

| REAGENT or RESOURCE | SOURCE | IDENTIFIER |
|---|---|---|
| Antibodies | | |
| Rat monoclonal anti-HA | Sigma | Cat# 11867431001; RRID: AB_390919 |
| Rabbit anti-Parvalbumin | Swant | Cat# PV27; RRID: AB_2631173 |
| Mouse monoclonal anti-Glutamate Decarboxylase 67 | Millipore-Merk | Cat# G5419; RRID: AB_261978 |
| Goat anti-rat Alexa Fluor 488 | Invitrogen | Cat# A-11006; RRID: AB_2534074 |
| Goat anti-rabbit Cy5 | Jackson ImmunoResearch | Cat# 711-175-152; RRID: AB_2340607 |
| Donkey anti-rabbit Cy3 | Jackson ImmunoResearch | Cat# 711-165-152; RRID: AB_2307443 |
| Goat anti-mouse Alexa Fluor 647 | Jackson ImmunoResearch | Cat# 115-605-003; RRID: AB_2338902 |
| DNA and Plasmids | | |
| C4b mouse sequence | NCBI Resources | NM_009780.2 |
| pCAG-HAC4b | This paper | N/A |
| pCAG-TdTomato | This paper | N/A |
| Experimental Models: Organisms/Strains | | |
| Mouse: C57Bl6/J | Janvier Labs | C57BL/6JRj |
| Mouse: PV-Cre (B6;129P2-Pvalb$^{tm1(cre)Arbr/J}$) | The Jackson Laboratory | JAX: 017320 |
| Mouse: ChR2-YFP-loxP (B6;129S-Gt(ROSA)$^{26Sortm32(CAG-COP4*H134R/EYFP)Hze/J}$) | The Jackson Laboratory | JAX: 012569 |
| Mouse: RCE-loxP (Gt(ROSA)$^{26Sortm1.1(CAG-EGFP)Fsh/Mmjax}$) | The Jackson Laboratory | JAX : 032037 |
| Software and Algorithms | | |
| GraphPad Prism | https://www.graphpad.com/ | RRID: SCR_002798 |
| ImageJ | https://imagej.nih.gov/ij/ | RRID: SCR_003070 |
| MiniAnalysis | http://www.synaptosoft.com/MiniAnalysis/ | RRID: SCR_002184 |
| IgorPro | https://www.wavemetrics.com/ | RRID: SCR_000325 |
| PatchMaster | http://www.heka.com/products/products_main.html#soft_pm | RRID:SCR_000034 |
| NeuroMatic | http://www.neuromatic.thinkrandom.com/ | RRID:SCR_004186 |
| Imetronic | http://www.imetronic.com/ | N/A |
| Viewpoint | https://viewpoint.fr/en/p/software/videotrack | N/A |

### Supplementary figure and table legends

##### Supplementary Figure S1. C4 overexpression in prefrontal cortex using *in utero* electroporation

***(A)*** Schematic drawing illustrating the *in utero* electroporation (IUE) method at embryonic day 15 (E15) for specific targeting of layer II/III PFC pyramidal neurons. VZ: ventricular zone.

***(B)*** Plasmid constructs used for IUE. Embryonic brains were co-electroporated with pCAG-HA-C4 and pCAG-tdTomato for C4 overexpression (C4-OE), or only with pCAG-tdTomato (control). pCAG-HA-C4 and pCAG-tdTomato were co-electroporated with a 4:1 ratio to ensure that most tdTomato+ cells overexpressed C4.

***(C)*** Left panel: Electroporated brain at postnatal day 30 (P30) exhibiting TdTomato expression in PFC; scale bar: 5 mm. Middle panel: Sagittal slice showing the distribution of TdTomato positive cells in PCF layer II/III. OB: olfactive bulb; scale bar: 500 μm. Right panel: typical morphology of an electroporated pyramidal cell; scale bar: 30 μm.

***(D)*** Immunostaining for TdTomato (red) and HA (green) in PFC layer II/III at P30. The three panels on the right are enlargements of the boxed area in the left panel. Note that tdTomato+ neurons also expressed HA-C4; scale bar: 100 µm; scale bar in magnified image: 20 µm.

***(E)*** Quantification of the results exemplified in (D).

**Supplementary Figure S2. Similar laminar position of electroporated pyramidal cells in control and C4-OE mice**

Left: sagittal PFC sections from control and C4-OE mice; scale bar: 300µm. Right, enlargements of the boxed area in the left panel showing PFC layers. All tdTomato+ cells are located in PFC layer 3; scale bar: 100 µm; M2: motor cortex; PFC: prefrontal cortex; OB: olfactory bulb.

**Supplementary Figure S3. Passive and active neuronal properties in control and C4-OE mice**

***(A)*** Sample spike trains evoked by a 400 pA somatic current injection in a control and in a C4-OE pyramidal cell; scale bars: 20 mV/100 ms.

***(B)*** f-i curves for control (n = 20 cells from 3 mice) and C4-OE neurons (n = 17 cells from 3 mice).

***(C)*** Input resistance (left), Cell capacitance (middle) and resting membrane potential (right) in control (n = 20 cells from 3 mice) and C4-OE neurons (n = 17 cells from 3 mice). Neither firing frequency nor intrinsic neuronal properties differed between control and C4-OE pyramidal cells. All data are represented as mean ± SEM. Each open circle represents an individual neuron.

**Supplementary Figure S4. Light-evoked firing of ChR2-expressing PV neurons**

***(A)*** Homozygous PV-Cre::RCE females were crossed with homozygous lox-STOP-lox ChR2-YFP males before electroporation of timed-pregnant females with tdTomato or C4 + tdTomato. Electroporated offspring (age P25-P30) were used for ex vivo patch-clamp and optogenetics.

***(B)*** Left, patch-clamped YFP+ cell (left); right, spike train evoked by a 500 ms pulse of blue (470 nm) light. YFP+ cells exhibited the typical fast-spiking pattern of PV neurons (n=3).

***(C)*** Left, light-evoked IPSCs were recorded at a holding potential of +10 mV in electroporated PFC pyramidal cells from control or C4-OE PVCre-RCE::lox-STOP-lox ChR2-YFP mice. Right, light-evoked IPSC blocked by the addition of of Gabazine (10 μM), demonstrating its GABAergic nature; scale bars: 50 pA/ 50ms.

**Supplementary Figure S5. Laminar distribution of PV neurons in control and C4-OE mice**

***(A)*** PV staining in PFC sagittal slices from a control and from a C4-OE mouse; scale bar: 100 µm.

***(B)*** The repartition by layer (left) as well as the total number of histologically identified PV (right) is unchanged in C4-OE (n = 10 mice) compared to control (n = 9 mice) mice. All data are represented as mean ± SEM.

**Supplementary Figure S6. Increased excitatory inputs onto on PV neurons in C4-OE mice**

***(A)*** Sample traces of mEPSCs recordings in PV interneurons from control and C4-OE juvenile (P25-P30) mice recorded at −65 mV in the presence of TTX and GABAzine; scale bars: 20 pA/100ms.

***(B)*** mEPSC frequency, but not IPSC amplitude, was lower in C4-OE mice than in control mice (frequency: Mann-Whitney test, * $P = 0.0184$).

***(C)*** Average of 12 consecutive recordings; EPSCs were evoked by paired-pulse stimulation and recorded in PV interneurons from control and C4-OE mice at a holding potential of -65 mV; scale bars: 50 pA/50 ms.

***(D)*** Summary histogram of the paired-pulse ratio in control and C4-OE.

***(E)*** Sample traces of mIPSCs recorded at a holding potential of −65 mV with a high-chloride intracellular solution in PV interneurons from control and C4-OE juvenile (P25-P30) mice, in the presence of TTX, APV and CNQX; scale bar: 20 pA/100 ms.

***(F)*** Mean frequency (left) and amplitude (right) of mIPSC in control and C4-OE neurons.

All data are represented as mean ± SEM. Open circles represent single cells.

**Supplementary Table S1. Summary table of patch-clamp results (pyramidal cells)**

N refers to the number of mice, n refers to the number of neuronal recordings. All data are presented as mean ± SEM. Sidak's test # $P < 0.05$; ## $P < 0.01$; ### $P < 0.001$; #### $P < 0.0001$

**Supplementary Table S2. Summary table of imaging results**

N refers to the number of mice, n refers to the number of dendrites or ROI as specified in each experiments. All data are presented as mean ± SEM.

**Supplementary Table S3. Summary table of patch-clamp results (PV interneurons)**

N refers to the number of mice, n refers to the number of neuronal recordings. All data are presented as mean ± SEM.

**Supplementary Table S4. Summary table of behavioral studies**

N refers to the number of mice. All data are presented as mean ± SEM.

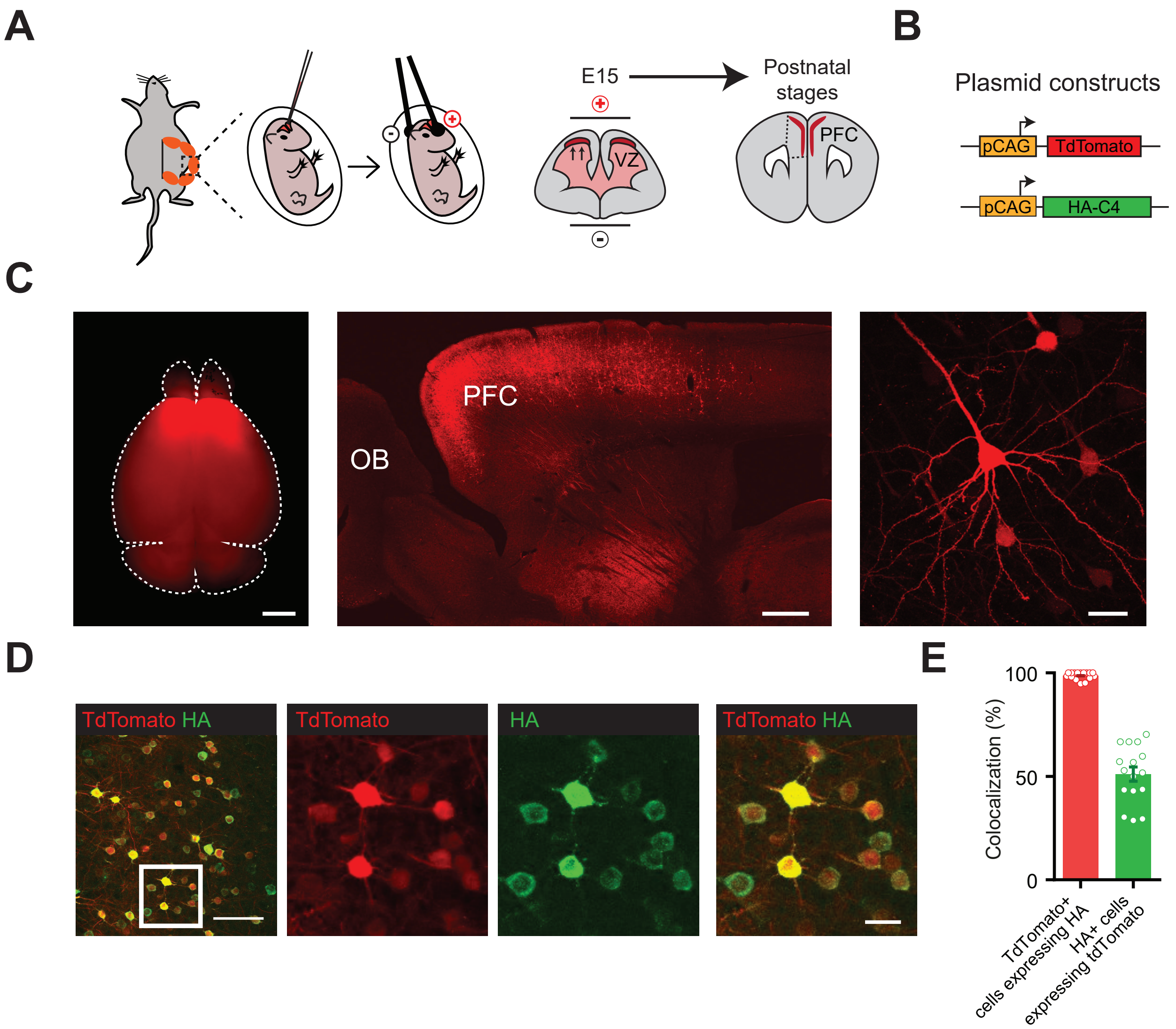

**Supplementary Figure 1**

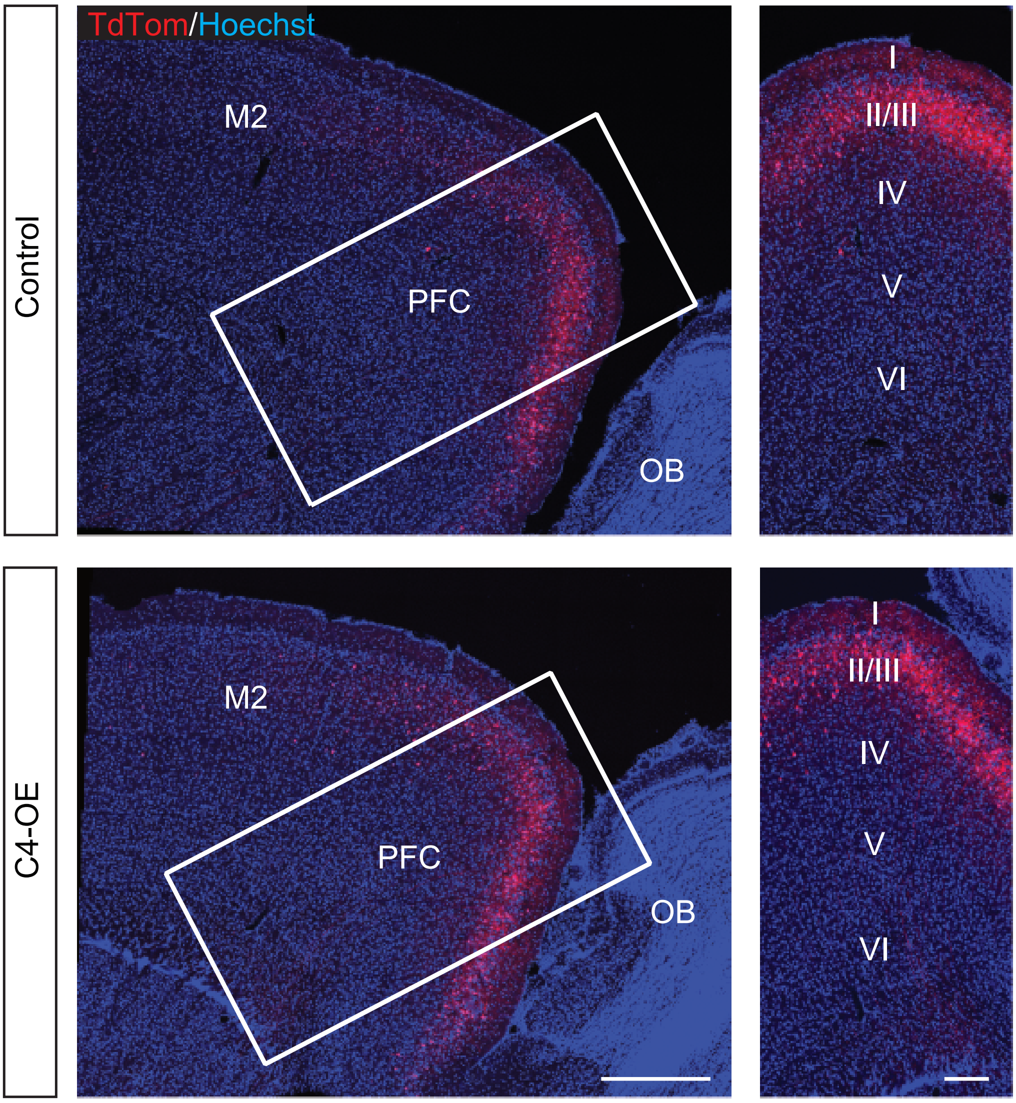

**Supplementary Figure 2**

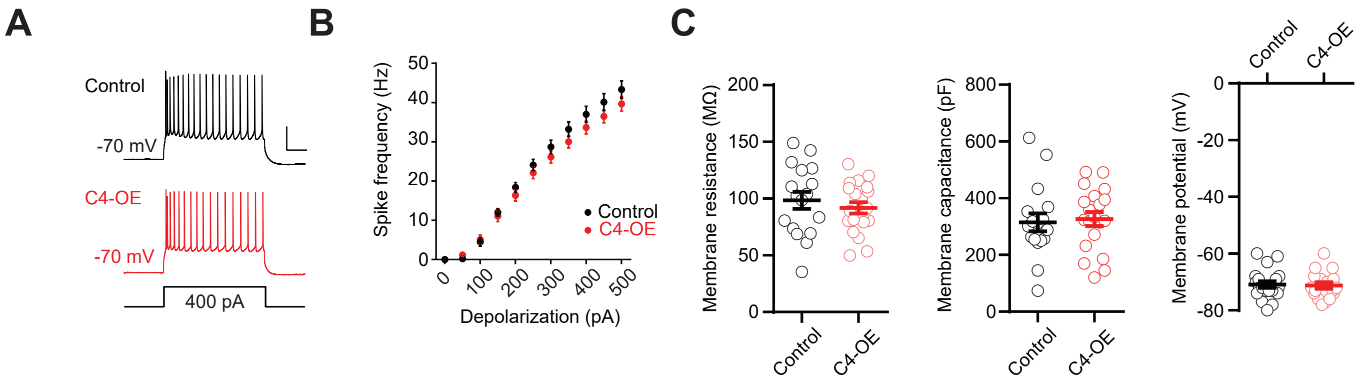

**Supplementary Figure 3**

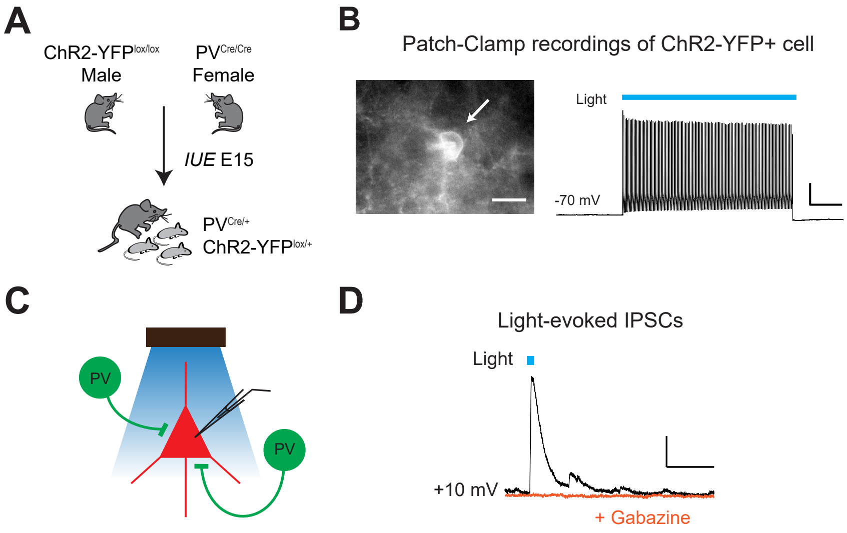

#### Supplementary Figure 4

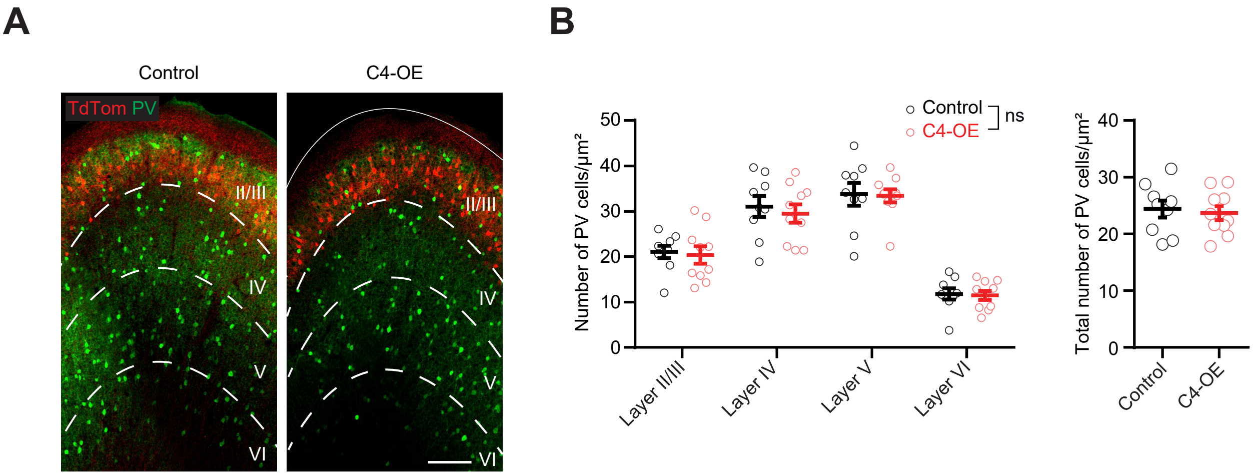

**Supplementary Figure 5**

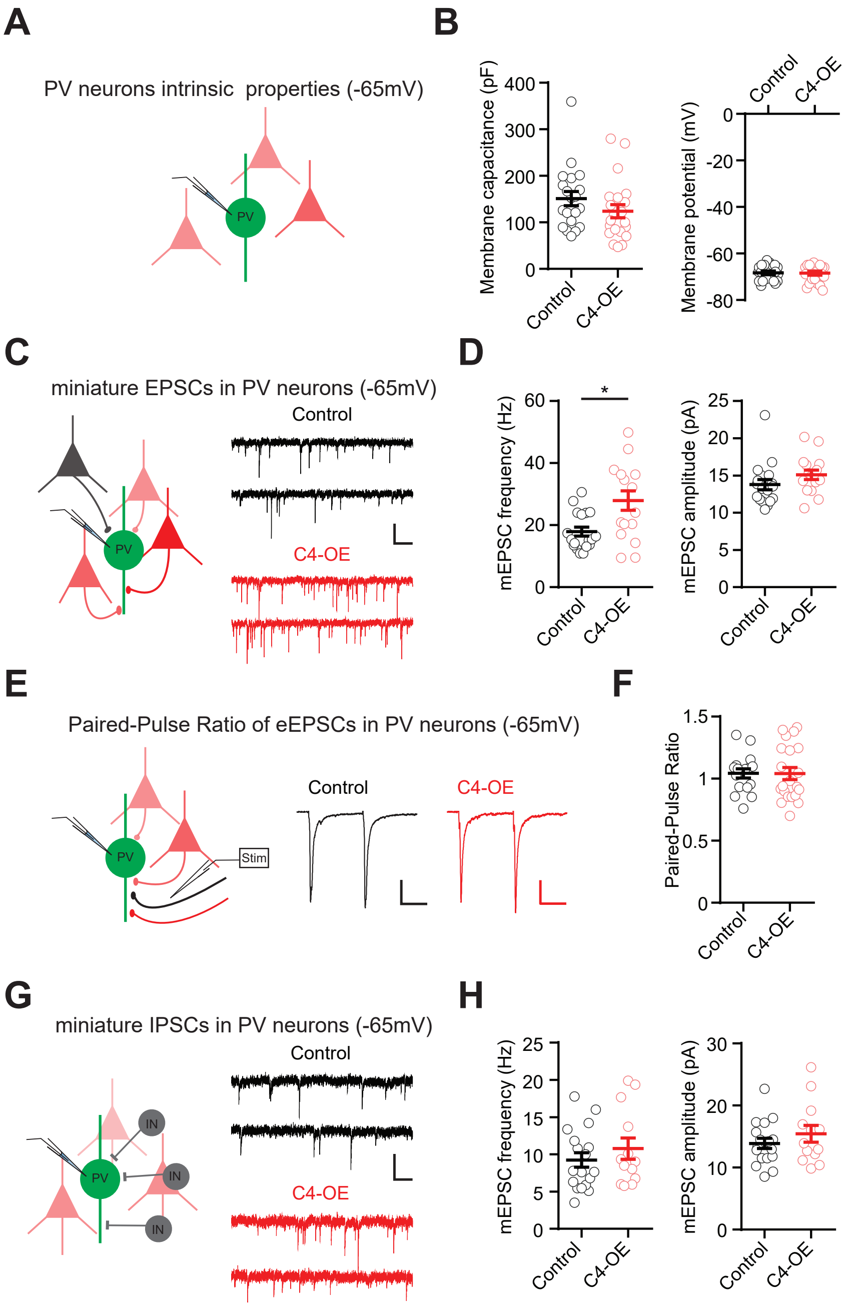

**Supplementary Figure 6**

#### Table S1

| | Control | n | N | C4-OE | n | N | Test | p value | Significant | F statistics (ANOVA) |
|---|---|---|---|---|---|---|---|---|---|---|
| **Intrinsic properties at P25-P30** | | | | | | | | | | |
| Membrane resistance (MΩ) | 91.9 ± 4.97 | 20 | 3 | 98.6 ± 7.62 | 17 | 3 | T-test | 0.456 | ns | |
| Resting membrane potential (mV) | -70.9 ± 1.16 | 20 | 3 | -71.2 ± 1.17 | 17 | 3 | T-test | 0.841 | ns | |
| Membrane capacitance (pF) | 327 ± 24.7 | 20 | 3 | 315 ± 31.6 | 17 | 3 | T-test | 0.773 | ns | |
| Firing frequency | | 20 | 4 | | 17 | 3 | Two-way RM ANOVA | 0.304 | ns | $F_{(1,35)} = 1.09$ |
| **mEPSC frequency (Hz)** | | | | | | | Two-way ANOVA | C4 factor: <0.0001 | **** | $F_{(1,105)} = 17.4$ |
| | | | | | | | | Age factor: <0.0001 | **** | $F_{(2,105)} = 54.4$ |
| | | | | | | | | Interaction: 0.0059 | ** | $F_{(2,105)} = 5.4$ |
| *P9-P10* | 2.03 ± 0.17 | 21 | 3 | 2.02 ± 0.15 | 18 | 4 | Sidak's post-hoc test | | ns | |
| *P25-P30* | 10.2 ± 1.31 | 16 | 5 | 6.20 ± 0.90 | 14 | 6 | Sidak's post-hoc test | | #### | |
| *P150-P220* | 4.58 ± 0.36 | 20 | 6 | 2.59 ± 0.21 | 22 | 5 | Sidak's post-hoc test | | # | |
| *Control: P9-P10 vs P25-P30* | | | | | | | Sidak's post-hoc test | | #### | |
| *Control: P25-P30 vs P150-P220* | | | | | | | Sidak's post-hoc test | | #### | |
| *Control: P9-P10 vs P150-P220* | | | | | | | Sidak's post-hoc test | | ## | |
| *C4-OE: P9-P10 vs P25-P30* | | | | | | | Sidak's post-hoc test | | #### | |
| *C4-OE: P25-P30 vs P150-P220* | | | | | | | Sidak's post-hoc test | | ### | |
| *C4-OE: P9-P10 vs P150-P220* | | | | | | | Sidak's post-hoc test | | ns | |
| **mEPSC amplitude (pA)** | | | | | | | Two-way ANOVA | C4 factor: 0.399 | ns | $F_{(1,105)} = 0.716$ |
| | | | | | | | | Age factor: <0.0001 | **** | $F_{(2,105)} = 85.82$ |
| | | | | | | | | Interaction: 0.477 | ns | $F_{(2,105)} = 0.745$ |
| *P9-P10* | 17.2 ± 1.241 | 21 | 3 | 17.7 ± 0.855 | 18 | 4 | Sidak's post-hoc test | | ns | |
| *P25-P30* | 8.87 ± 0.586 | 16 | 5 | 8.10 ± 0.478 | 14 | 6 | Sidak's post-hoc test | | ns | |
| *P150-P220* | 9.927 ± 0.442 | 20 | 6 | 8.637 ± 0.350 | 22 | 5 | Sidak's post-hoc test | | ns | |
| *Control: P9-P10 vs P25-P30* | | | | | | | Sidak's post-hoc test | | #### | |
| *Control: P25-P30 vs P150-P220* | | | | | | | Sidak's post-hoc test | | ns | |
| *Control: P9-P10 vs P150-P220* | | | | | | | Sidak's post-hoc test | | #### | |
| *C4-OE: P9-P10 vs P25-P30* | | | | | | | Sidak's post-hoc test | | #### | |
| *C4-OE: P25-P30 vs P150-P220* | | | | | | | Sidak's post-hoc test | | ns | |
| *C4-OE: P9-P10 vs P150-P220* | | | | | | | Sidak's post-hoc test | | #### | |
| **Excitatory PPR** | | | | | | | Two-way ANOVA | C4 factor: 0.809 | ns | $F_{(1,105)} = 0.033$ |
| | | | | | | | | Age factor: 0.0011 | ** | $F_{(1,105)} = 7.31$ |
| | | | | | | | | Interaction: 0.806 | ns | $F_{(1,105)} = 0.213$ |
| *P9-P10* | 0.916 ± 0.056 | 16 | 3 | 0.875 ± 0.031 | 21 | 3 | Sidak's post-hoc test | | ns | |
| *P25-P30* | 1.04 ± 0.04 | 17 | 3 | 1.04 ± 0.05 | 21 | 3 | Sidak's post-hoc test | | ns | |

| | | | | | | | | | |
|---|---|---|---|---|---|---|---|---|---|
| *P150-P220* | 1.06 ± 0.07 | 19 | 5 | 1.08 ± 0.04 | 17 | 4 | Sidak's post-hoc test | | ns | |
| *Control: P9-P10 vs P25-P30* | | | | | | | Sidak's post-hoc test | | ns | |
| *Control: P25-P30 vs P150-P220* | | | | | | | Sidak's post-hoc test | | ns | |
| *Control: P9-P10 vs P150-P220* | | | | | | | Sidak's post-hoc test | | ns | |
| *C4-OE: P9-P10 vs P25-P30* | | | | | | | Sidak's post-hoc test | | ns | |
| *C4-OE: P25-P30 vs P150-P220* | | | | | | | Sidak's post-hoc test | | ns | |
| *C4-OE: P9-P10 vs P150-P220* | | | | | | | Sidak's post-hoc test | | # | |
| **AMPA/NMDA ratio** | | | | | | | Two-way ANOVA | C4 factor: <0.0001 | **** | $F_{(1, 108)} = 18.9$ |
| | | | | | | | | Age factor: <0.0001 | **** | $F_{(2, 108)} = 40.6$ |
| | | | | | | | | Interaction: 0.007 | ** | $F_{(1, 108)} = 5.2$ |
| *P9-P10* | 0.856 ± 0.078 | 17 | 3 | 0.844 ± 0.085 | 21 | 3 | Sidak's post-hoc test | | ns | |
| *P25-P30* | 0.928 ± 0.028 | 23 | 4 | 1.294 ± 0.086 | 20 | 5 | Sidak's post-hoc test | | ### | |
| *P150-P220* | 1.317 ± 0.073 | 17 | 4 | 1.752 ± 0.085 | 16 | 4 | Sidak's post-hoc test | | ### | |
| *Control: P9-P10 vs P25-P30* | | | | | | | Sidak's post-hoc test | | ns | |
| *Control: P25-P30 vs P150-P220* | | | | | | | Sidak's post-hoc test | | ### | |
| *Control: P9-P10 vs P150-P220* | | | | | | | Sidak's post-hoc test | | ### | |
| *C4-OE: P9-P10 vs P25-P30* | | | | | | | Sidak's post-hoc test | | #### | |
| *C4-OE: P25-P30 vs P150-P220* | | | | | | | Sidak's post-hoc test | | ### | |
| *C4-OE: P9-P10 vs P150-P220* | | | | | | | Sidak's post-hoc test | | #### | |
| **NMDA decay time (ms)** | | | | | | | Two-way ANOVA | C4 factor: 0.490 | ns | $F_{(1, 108)} = 0.48$ |
| | | | | | | | | Age factor: <0.0001 | **** | $F_{(2, 108)} = 73.09$ |
| | | | | | | | | Interaction: 0.332 | ns | $F_{(1, 108)} = 1.12$ |
| *P9-P10* | 0.098 ± 0.008 | 17 | 3 | 0.104 ± 0.006 | 21 | 3 | Sidak's post-hoc test | | ns | |
| *P25-P30* | 0.056 ± 0.003 | 23 | 4 | 0.051 ± 0.002 | 20 | 5 | Sidak's post-hoc test | | ns | |
| *P150-P220* | 0.058 ± 0.003 | 17 | 4 | 0.050 ± 0.002 | 16 | 4 | Sidak's post-hoc test | | ns | |
| *Control: P9-P10 vs P25-P30* | | | | | | | Sidak's post-hoc test | | #### | |
| *Control: P25-P30 vs P150-P220* | | | | | | | Sidak's post-hoc test | | ns | |
| *Control: P9-P10 vs P150-P220* | | | | | | | Sidak's post-hoc test | | #### | |
| *C4-OE: P9-P10 vs P25-P30* | | | | | | | Sidak's post-hoc test | | #### | |
| *C4-OE: P25-P30 vs P150-P220* | | | | | | | Sidak's post-hoc test | | ns | |
| *C4-OE: P9-P10 vs P150-P220* | | | | | | | Sidak's post-hoc test | | #### | |
| **Rectification Index** | | | | | | | Two-way ANOVA | C4 factor: <0.0001 | **** | $F_{(1, 112)} = 18.09$ |
| | | | | | | | | Age factor: 0.0058 | ** | $F_{(2, 112)} = 5.40$ |
| | | | | | | | | Interaction: 0.196 | ns | $F_{(2,112)} = 1.65$ |
| *P9-P10* | 1.09 ± 0.06 | 20 | 3 | 1.02 ± 0.05 | 20 | 3 | Sidak's post-hoc test | | ns | |
| *P25-P30* | 1.31 ± 0.05 | 21 | 3 | 1.06 ± 0.05 | 16 | 3 | Sidak's post-hoc test | | ## | |

| | | | | | | | | | |
|---|---|---|---|---|---|---|---|---|---|
| *P150-P220* | 1.13 ± 0.03 | 17 | 4 | 0.97 ± 0.03 | 24 | 3 | Sidak's post-hoc test | | # | |
| *Control: P9-P10 vs P25-P30* | | | | | | | Sidak's post-hoc test | | ## | |
| *Control: P25-P30 vs P150-P220* | | | | | | | Sidak's post-hoc test | | # | |
| *Control: P9-P10 vs P150-P220* | | | | | | | Sidak's post-hoc test | | ns | |
| *C4-OE: P9-P10 vs P25-P30* | | | | | | | Sidak's post-hoc test | | ns | |
| *C4-OE: P25-P30 vs P150-P220* | | | | | | | Sidak's post-hoc test | | ns | |
| *C4-OE: P9-P10 vs P150-P220* | | | | | | | Sidak's post-hoc test | | ns | |
| **mIPSC Amplitude at P25-P30 (pA)** | 17.1 ± 0.98 | 16 | 4 | 12.74 ± 0.46 | 16 | 6 | T-test | 0.0003 | *** | |
| **mIPSC Frequency at P25-P30 (Hz)** | 7.08 ± 0.57 | 16 | 4 | 4.46 ± 0.61 | 16 | 6 | Mann-Whitney test | 0.0004 | *** | |
| **Short-term plasticity of evoked IPSCs at P25-P30 (electrical stim)** | | 29 | 3 | | 37 | 3 | Two-way RM ANOVA | 0.089 | ns | $F_{(1,64)} = 2.988$ |
| *Pulse II/ pulse I* | 0.542 ± 0.028 | 29 | 3 | 0.610 ± 0.027 | 37 | 3 | Sidak's post-hoc test | | ns | |
| *Pulse III/ pulse I* | 0.437 ± 0.025 | 29 | 3 | 0.488 ± 0.024 | 37 | 3 | Sidak's post-hoc test | | ns | |
| *Pulse IV/ pulse I* | 0.397 ± 0.022 | 29 | 3 | 0.442 ± 0.021 | 37 | 3 | Sidak's post-hoc test | | ns | |
| **Short-term plasticity of evoked IPSCs at P25-P30 (optogenetics)** | | 43 | 4 | | 30 | 3 | Two-way RM ANOVA | 0.014 | * | $F_{(1,71)} = 6.85$ |
| *Pulse II/ pulse I* | 0.541 ± 0.016 | 43 | 4 | 0.592 ± 0.021 | 30 | 3 | Sidak's post-hoc test | | ns | |
| *Pulse III/ pulse I* | 0.400 ± 0.013 | 43 | 4 | 0.465 ± 0.019 | 30 | 3 | Sidak's post-hoc test | | # | |
| *Pulse IV/ pulse I* | 0.357 ± 0.014 | 43 | 4 | 0.411 ± 0.018 | 30 | 3 | Sidak's post-hoc test | | ns | |

**Table S2**

| | Control | n | N | C4-OE | n | N | Test | p value | Significant | F statistics (ANOVA) |
|---|---|---|---|---|---|---|---|---|---|---|
| ***In vivo* 2-photon imaging at P30 (n = dendrites ; N = mice)** | | | | | | | | | | |
| Spine density | 0.880 ± 0.017 | 99 | 10 | 0.672 ± 0.018 | 76 | 7 | T-test | < 0.0001 | **** | |
| ***In vivo* 2-photon imaging at P15-P18 (n = dendrites ; N = mice)** | | | | | | | | | | |
| Spine density at 0h | 1.15 ± 0.03 | 73 | 5 | 0.927 ± 0.025 | 55 | 7 | T-test | < 0.0001 | **** | |
| Spine density from 0h to 4h | | | 5 | | | 7 | Two-way RM ANOVA | C4 factor: < 0.0001 | **** | $F_{(1,10)} = 44.8$ |
| | | | | | | | | Time factor: 0.780 | ns | $F_{(1,10)} = 0.083$ |
| | | | | | | | | Time factor: 0.366 | ns | $F_{(1,10)} = 0.899$ |
| Stable spine density | 1.05 ± 0.02 | 73 | 5 | 0.89 ± 0.03 | 55 | 7 | T-test | < 0.0001 | **** | |
| Gained spine density | 0.092 ± 0.008 | 73 | 5 | 0.047 ± 0.004 | 55 | 7 | T-test | < 0.0001 | **** | |
| Lost spine density | 0.095 ± 0.005 | 73 | 5 | 0.039 ± 0.006 | 55 | 7 | Mann-Whitney test | < 0.0001 | **** | |
| Turnover rate (%) | 8.81 ± 0.470 | 73 | 5 | 5.26 ± 0.53 | 55 | 7 | Mann-Whitney test | < 0.0001 | **** | |
| **GAD 67 expression (n = ROI ; N = mice)** | | | | | | | | | | |
| Frequency distribution | | 117 | 6 | | 134 | 6 | Kolmogorov-Smirnov Test | < 0.0001 | **** | |
| Mean intensity | 28.37 ± 0.815 | 117 | 6 | 25.46 ± 0.54 | 134 | 6 | Mann-Whitney test | 0.0114 | * | |
| **PV cell distribution** | | | 9 | | | 10 | Two-way RM ANOVA | 0.705 | ns | $F_{(1,17)} = 0.148$ |
| *Layer II/III* | 21.1 ± 1.4 | | 9 | 20.4 ± 1.9 | | 10 | Sidak's post-hoc test | | ns | |
| *Layer IV* | 31.1 ± 2.3 | | 9 | 29.5 ± 2.0 | | 10 | Sidak's post-hoc test | | ns | |
| *Layer V* | 33.8 ± 2.5 | | 9 | 33.4 ± 1.5 | | 10 | Sidak's post-hoc test | | ns | |
| *Layer VI* | 11.8 ± 1.2 | | 9 | 11.5 ± 1.0 | | 10 | Sidak's post-hoc test | | ns | |
| **PV cell total number** | 24.4 ± 1.5 | | 9 | 23.7 ± 1.21 | | 10 | T-test | 0.705 | ns | |

**Table S3**

| | Control | n | N | C4-OE | n | N | Test | p value | Significant | F statistics (ANOVA) |
|---|---|---|---|---|---|---|---|---|---|---|
| **Intrinsic properties** | | | | | | | | | | |
| Membrane resistance (MΩ) | 129.4 ± 8.8 | 20 | 3 | 98.3 ± 7.0 | 21 | 4 | Mann-Whitney test | 0.0024 | ** | |
| Resting membrane potential (mV) | -68.0 ± 0.8 | 20 | 3 | -68.6 ± 0.8 | 21 | 4 | T-test | 0.881 | ns | |
| Membrane capacitance (pF) | 151 ± 15 | 20 | 3 | 124 ± 14 | 21 | 4 | Mann-Whitney test | 0.127 | ns | |
| Firing | | 20 | 3 | | 21 | 4 | Two-way RM ANOVA | 0.013 | * | $F_{(1,37)} = 6.82$ |
| **mEPSC Amplitude (pA)** | 13.8 ± 0.7 | 18 | 5 | 15.1 ± 0.6 | 16 | 6 | Mann-Whitney test | 0.069 | ns | |
| **mEPSC Frequency (Hz)** | 17.9 ± 1.4 | 18 | 5 | 27.9 ± 3.1 | 16 | 6 | Mann-Whitney test | 0.018 | * | |
| **Excitatory PPR** | 1.04 ± 0.06 | 24 | 4 | 1.12 ± 0.05 | 21 | 3 | T-test | 0.251 | ns | |
| **mIPSC Amplitude (pA)** | 13.9 ± 0.8 | 18 | 4 | 15.4 ± 1.4 | 13 | 4 | T-test | 0.312 | ns | |
| **mIPSC Frequency (Hz)** | 9.25 ± 0.97 | 18 | 4 | 10.8 ± 1.43 | 13 | 4 | T-test | 0.365 | ns | |

#### Table S4

| | Control | N | C4-OE | N | Test | p value | Significant | F statistics (ANOVA) |
|---|---|---|---|---|---|---|---|---|
| **Locomotion** | | | | | | | | |
| Actimeter (repeated measures) | | 17 | | 13 | Two-way RM ANOVA | 0.416 | ns | $F_{(1,28)} = 0.681$ |
| Actimeter total activity | 353 ± 30 | 17 | 404 ± 59 | 13 | Mann-Whitney test | 0.920 | ns | |
| Open field (repeated measures) | | 5 | | 7 | Two-way RM ANOVA | 0.613 | ns | $F_{(1,90)} = 0.258$ |
| Open field total activity | 1765 ± 15 | 5 | 1849 ± 230 | 7 | Mann-Whitney test | 0.86 | ns | |
| **Working memory** | | | | | | | | |
| Non-matching to sample | | 15 | | 14 | Two-way RM ANOVA | 0.556 | ns | $F_{(1,135)} = 0.348$ |
| Odor Span | 8.40 ± 0.83 | 15 | 4.43 ± 0.54 | 14 | Mann-Whitney test | 0.0022 | ** | |
| Spatial Span | 6.40 ± 0.73 | 15 | 4.21 ± 0.68 | 14 | Mann-Whitney test | 0.045 | * | |
